# *In vivo* high-throughput screening of novel adeno-associated viral capsids targeting adult neural stem cells in the subventricular zone

**DOI:** 10.1101/2021.03.05.434064

**Authors:** Sascha Dehler, Lukas PM Kremer, Santiago Cerrizuela, Thomas Stiehl, Jonas Weinmann, Heike Abendroth, Susanne Kleber, Alexander Laure, Jihad El Andari, Simon Anders, Anna Marciniak-Czochra, Dirk Grimm, Ana Martin-Villalba

**Author notes:** Contributed equally (Sascha Dehler, Lukas PM Kremer). Contributed equally (Santiago Cerrizuela, Thomas Stiehl). Correspondence: Prof. Dr. Ana Martin-Villalba.

## Abstract

The adult mammalian brain entails a reservoir of neural stem cells (NSCs) generating glial cells and neurons. However, NSCs become increasingly quiescent with age, which hampers their regenerative capacity. New means are therefore required to genetically modify adult NSCs for re-enabling endogenous brain repair. Recombinant adeno-associated viruses (AAVs) are ideal gene therapy vectors due to an excellent safety profile and high transduction efficiency. We thus conducted a high-throughput screening of 177 intraventricularly injected barcoded AAV variants profiled by RNA sequencing. Quantification of barcoded AAV mRNAs identified two synthetic capsids, AAV9_A2 and AAV1_P5, both of which transduce active and quiescent NSCs. Further optimization of AAV1_P5 by judicious selection of promoter and dose of injected viral genomes enabled labeling of 30-60% of the NSC compartment, which was validated by FACS analyses and single cell RNA sequencing. Importantly, transduced NSC readily produced neurons. The present study identifies AAV variants with a high regional tropism towards the v-SVZ with high efficiency in targeting adult NSCs, thereby paving the way for preclinical testing of regenerative gene therapy.

## Introduction

The adult brain has long been considered as a tissue with no regenerative capacity partly due to the absence of pluripotent cells. In the late 1990’s, a reservoir of neural stem cells (NSCs) with the potential to generate glia and neuronal progeny was identified in the adult mammalian brain^1,3^. The largest reservoir of NSCs in rodents is located along the walls of the lateral ventricles, the so-called ventricular-subventricular zone (v-SVZ). The potential of these NSCs to produce different glia and neuronal subtypes has been demonstrated by lineage-tracing studies^4–6^. NSCs get activated to provide progeny for tissue homeostasis but also in the frame of a traumatic brain injury^7–12^. However, the ability to activate NSCs highly declines with age^2^, hampering repair of the brain. This fairly limited endogenous regenerative capacity calls for new strategies to specifically target and genetically modify adult NSCs within the natural environment of the brain.

Many different viral and transgenic approaches have been developed in the past to manipulate adult NSCs and their progeny^13^. For a long time, onco-retroviruses and lentiviruses that integrate their genomes into the host cellular chromatin were the tools of choice. However, limitations of integrating viruses^14^, such as insertional mutagenesis^15,16^, gradual silencing of the inserted transgene^17,18^ and the fact that not all non-dividing cells are equally transduced *in vivo^19^* hamper their use for targeting of especially quiescent (q)NSCs within the v-SVZ. Over the last few years, the non-enveloped adeno-associated viral (AAV) vectors have taken center stage as a gene delivery vehicle for human gene therapy with two gene therapeutic approaches that have gained regulatory approval for commercial use in patients: Glybera (uniQure) and Luxturna (Novartis) and with a large amount of AAV gene therapeutic strategies even in the CNS under clinical development, as reviewed in^20,21^.

AAVs are small virus particles, belonging to the dependoviruses within the parvoviridae family with a capsid diameter of ~22nm that is sterically limiting its genome to ~4.7kb^22^. The original AAV genome consists of only two genes, the *rep* and *cap* gene, which are organized in three open reading frames. The *cap* gene determines the structure of the AAV capsid, while the *rep* gene is involved in several processes ranging from transcription initiation to packaging of the AAV genome. For vector production these genes are commonly delivered in trans and thus can be easily modified^23–30^. Over the last decades, hundreds of AAV isolates were identified in various species, with an interestingly high homology regarding their capsid protein amino acid sequences, e.g. up to 99% for the primate isolate AAV1 compared to the human isolate AAV6^31^. Favorable safety profiles combined with the ability to mediate long-term transgene expression and to efficiently target many different human tissues are major assets that make AAVs a preferred technology^23,32–35^.

Nonetheless, specific targeting of NSCs in the v-SVZ has remained challenging to date. While the most efficient wild-type (wt) serotype, AAV9, shows high transduction efficiency upon intravenous and intracranial injection, it mainly targets neurons and astrocytes, but not NSCs^36–38^. Just recently, the power of structure-guided DNA shuffling was used to develop the newly engineered AAV variant SCH9. This new variant was able to target cells in the v-SVZ including NSCs^39^. However, to date, the usefulness of AAV vectors for transduction of stem cells remains debated, mainly based on conflicting reports concerning their transduction efficiency as reviewed^40^. The variable regions of the VP protein, which is encoded by the *cap* gene, are involved in receptor binding and antibody recognition and thus modifications thereof can be used to guide targeting of specific cell types. Engineering of the AAV capsid for optimization of organ, region or cell specificity can be achieved by methods such as random *cap* gene mutation, DNA family shuffling or peptide display, combined with *in vivo* selection^39,41–47^. Most recently, barcoding of double-stranded encapsidated DNA and next-generation sequencing (NGS) were shown to allow for high-throughput screening of AAV capsid libraries^48,49^. Here we apply these barcoded AAV-libraries by intracerebroventricular injection of the adult rodent brain. Using a combination of NGS, immunohistochemistry, flow cytometry and mathematical modeling we validate transduction of the NSCs within the v-SVZ and their neurogenic lineage by the novel AAV capsid AAV1_P5.

## Results

To identify AAV capsids able to transduce NSCs in the v-SVZ with the highest transduction efficiency possible, we performed an NGS-based high-throughput screening of 177 different barcoded AAV capsid variants. These AAV variants comprise 12 AAV wts, 94 newly generated peptide display mutants based on these wts and 71 chimeric capsids generated through DNA family shuffling. Among the synthetic capsids are 24 previously published benchmarks, the remaining ones were generated as described in the Methods section, Table S4 and in greater detail in ^50^. All AAV capsid variants were uniquely barcoded (with a 15 nt long random DNA sequence) and packaged into an AAV vector expressing a CMV promoter-controlled eYFP (enhanced yellow fluorescent protein) that harbors the barcode in its 3’UTR. A library comprising either 91 (library #1 from ^50^) or 157 (library #3 from ^50^) capsid variants was directly injected into the lateral ventricles of the adult mouse brain (10^10^ viral genomes in 2μl per mouse (vg), Fig 1a, Fig S2a).

**Figure 1:**
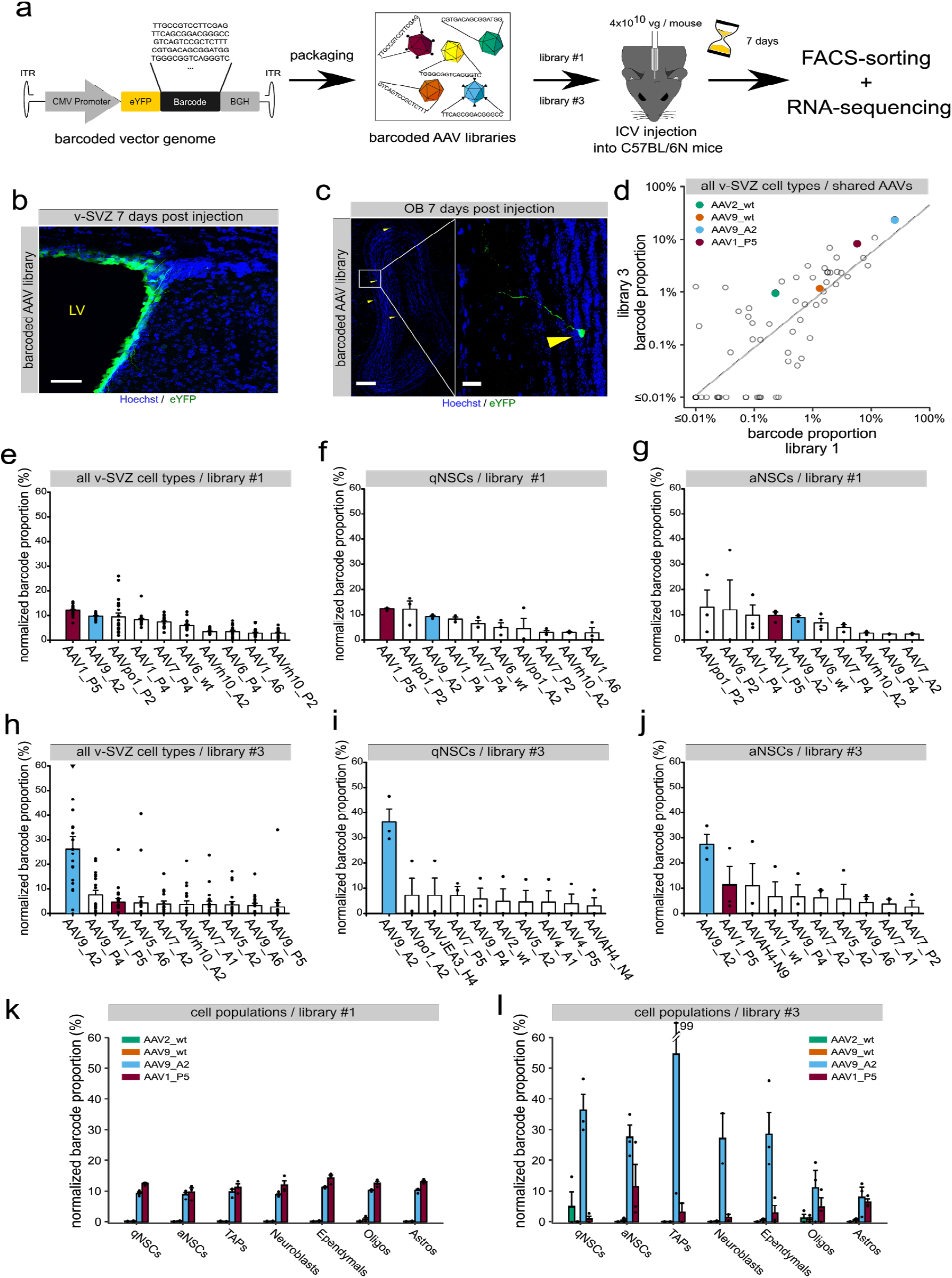
*In vivo* screening to identify AAV capsids that specifically target the v-SVZ. **a** Schematic illustration of the experimental outline to perform the *in vivo* screening. IHC of **b** the v-SVZ (scale bar 50 μm) or **c** the OB (scale bar 200 μm and 30 μm) after injection of library #1 into the lateral ventricle. **d** Mean barcode proportion over all FACS-sorted cell types for libraries #1 and #3. Only the 71 capsids shared between the two libraries are shown. **e** Barcode proportion in sample, adjusted for abundance in library (normalized barcode proportion) over all FACS-sorted cell types seven days after library #1 transduction; n=3 sets per cell type. **f,g** Normalized barcode read count seven days after library #1 transduction of **f** qNSCs or of **g** aNSCs; n=3 sets. **h** Normalized barcode read count over all FACS-sorted cell types seven days after library #3 transduction; n=2 sets for TAPs and neuroblasts, for all other cell types n=3 sets per cell type. **i,j** Normalized barcode read count seven days after library #3 transduction of **i** qNSCs or of **j** aNSCs; n=3 sets. **k,l** Normalized barcode read count of AAV2_wt, AAV9_wt, AAV9_A2 and AAV1_P5 after library #1 (**k**) and #3 (**l**) transduction of qNSCs, aNSCs, TAPs, neuroblasts, ependymal cells, astrocytes, and oligodendrocytes. All mice were eight weeks old at the time of AAV injection and all values are given as mean ± SEM. ITR, inverted terminal repeat; BGH, bovine growth hormone polyA signal; eYFP, enhanced yellow fluorescent protein; ICV, Intracerebroventricular. A set always consists of 6 mice. Three independent experiments were performed resulting in n = 3 sets (3 x 6 mice = 18 mice in total) (See also Figure S1 and S2).

7 days post-injection (dpi), quiescent and active NSCs (qNSCs or aNSCs, respectively), as well as other cell populations of the v-SVZ including transient amplifying progenitors (TAPs), neuroblasts, astrocytes, oligodendrocytes and ependymal cells, were FACS-sorted as previously described^2,7,51^ (Fig S1a-b, Tables S2 and S3). Finally, RNA libraries from the different cell populations were generated for NGS analysis (Fig 1a). In parallel, additional mice were sacrificed at 7 dpi for detection of the eYFP reporter in the v-SVZ. Efficient transduction of cells in the v-SVZ by both AAV libraries was confirmed by detecting the expression of the eYFP reporter along the ventricular walls (Fig 1b). Already after 7 dpi, few eYFP-positive (eYFP^+^) cells migrated to the olfactory bulb (OB) and were detected in the core and granular cell layer (GCL; Fig 1c), indicating that the AAV vector was retained along the lineage and did not prevent migration. For AAV mRNA analysis, capsids were ranked within each sorted cell population by the relative expression of their cognate barcodes, normalized by their frequency within library #1 and library #3. Overall capsid rankings of the 71 capsids shared by both libraries revealed the same top candidates and correlated strongly (Spearman’s rank correlation ρ = 0.84, p < 0.01) (Fig 1d). Furthermore, we did not find a significant association between barcode GC-content and frequency in either library (Fig S2l-m and Methods section), indicating that the results are not strongly influenced by GC-bias. Further analysis revealed that two synthetic capsids, AAV1_P5 and AAV9_A2 (peptide-modified derivatives of wt AAV1 or AAV9, respectively), stood out as the most efficient AAV capsid variants based on the ranking of their barcode enrichment (Fig 1d-j, Fig S2b-k). Notably, both active and quiescent NSCs were robustly transduced by these two AAV capsids (Fig 1f,g,i,j). Besides, AAV1_P5 and AAV9_A2 transduced other v-SVZ cell types, such as TAPs (Fig S2b,g), neuroblasts (Fig S2c,h), astrocytes (Fig S2d,i), oligodendrocytes (Fig S2e,j) and ependymal cells (Fig S2f,k). These two lead candidates clearly outperformed the well-established AAV2 and AAV9 wt capsids across all v-SVZ-cell populations (Fig 1k,l), as well as the parent wt AAV1. Taken together, our study has successfully identified AAV capsids that were highly region-specific for the v-SVZ, probably due to inability to migrate out of this region as reported for the SCH9 variant. These candidates exhibited a higher efficiency in targeting both active and quiescent NSCs than established wt AAV variants in the v-SVZ *in vivo.*

One potential application of gene therapy is to genetically modify freshly isolated cells and transplant them back to the donor. Hence, to identify the capsid with the fastest transduction rate of isolated NSCs, we assessed the expression dynamics of AAV2_wt, AAV9_wt, AAV9_A2 and AAV1_P5 in NSCs *in vitro.* To detect viral transduction of targeted cells and their progeny, we took advantage of the Cre/loxP system and engineered the AAVs to express a CAG promoter-controlled Cre recombinase fused to GFP (CAG_Cre::GFP). We decided to use the CAG promoter to assess performance of these capsids, since this promoter proved to outperform other promoters for *in utero* electroporation of embryonic neural progenitors^52^. Subsequently, we transduced primary cultured NSCs from tdTomato-flox mice (Fig 2a). Cre-fused GFP and cytoplasmic tdTomato were detected via immunocytochemistry at day 1, 3, 5 and 7 post-transduction (dpt) (Fig 2a-b). Interestingly, while all capsids showed a similar number of transduced cells at 7 dpt (Fig 2c-d), AAV1_P5 exhibited the fastest transduction kinetics (Fig 2c), already showing labeling at day 1 (Fig S3a,b).

**Figure 2:**
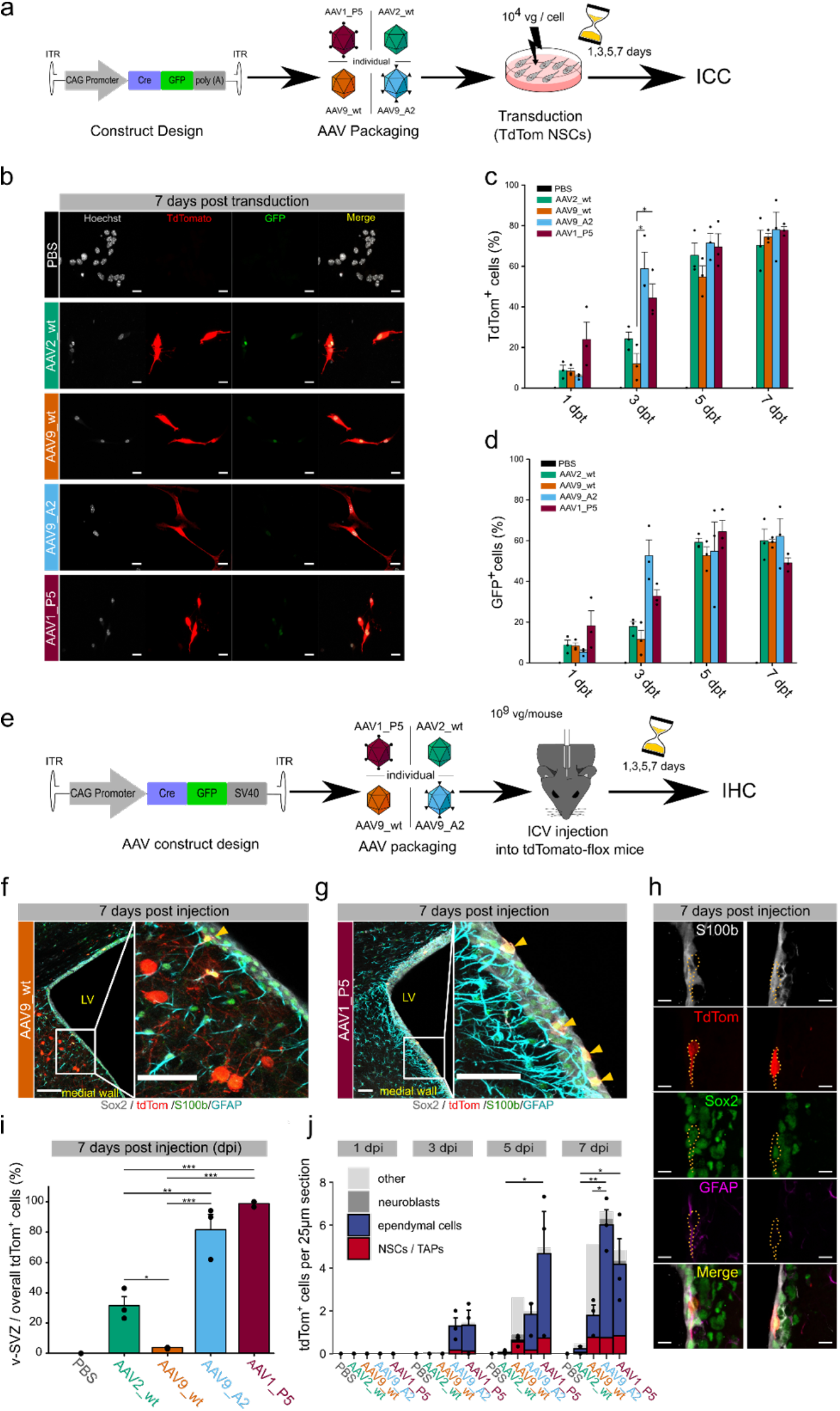
Assessment of expression dynamics and v-SVZ targeting of the lead candidate AAV capsids. **a** Experimental outline to assess expression dynamics of AAV1_P5, AAV9_A2 and two wt capsids *in vitro.* **b** Representative images of NSCs *in vitro* transduced with different AAV capsids 7 days after injection; scale bar 20 μm. **c** Dynamics of tdTomato expression at different time points in primary cultured NSCs. AAV9_wt_3dpt (11.9% ± 5.04) vs. AAV9_A2_3dpt (58.8% ± 8.24) vs. AAV1_3dpt (44.4% ± 6.94) (Kruskal-Wallis test followed by Dunn’s post-hoc test). **d** Dynamics of GFP expression at different time points in primary cultured NSCs. **c,d** Cultured NSCs were used up to passage 7, n=3 cell cultures from 3 different mice. **e** Schematic illustration of the experimental outline to *in vivo* validate different AAV capsids. **f,g** IHC of the v-SVZ with markers to discriminate the different cell types after **f** AAV9_wt and **g** AAV1_P5 transduction (scale bar 100 μm and 50 μm, respectively). **h** Markers for IHC used to discriminate the different cell types (NSCs left, ependymal cells right; scale bar 30 μm). **i** Proportion of tdTomato-labeled cells located in the v-SVZ among all tdTomato-positive cells in a 25 μm thick coronal brain section. A high proportion indicates regional specificity for the v-SVZ. AAV2_wt (31.5% ± 5.9) vs. AAV9_wt (3.84% ± 0.33) vs. AAV9_A2 (81.6% ±10.1) vs. AAV1_P5 (98.9% ± 1.13). **j** Dynamics of tdTomato expression at different time points in the full v-SVZ. Bars are partitioned by the mean proportion of cell types across mice. AAV2_wt_5dpi (0.06 ± 0.06) vs. AAV1_P5_5dpi (4.67 ± 1.96) and AAV2_wt_7dpi (0.22 ± 0.11) vs. AAV9_A2_7dpi (6.02 ± 0.71) vs. AAV1_P5_7dpi (4.17 ± 1.20). (See also Figure S3)

Next, we investigated whether the newly identified AAV capsids AAV1_P5 and AAV9_A2 also target v-SVZ cells *in vivo.* To this end, we individually injected 10^9^ vg of AAV9_A2, AAV1_P1 or the well-established wt serotypes AAV9 and AAV2, all containing the CAG_Cre::GFP construct, into tdTomato-flox mice (Fig 2e). Notably, at 7 dpi, the tropism towards the v-SVZ highly differed between the tested capsids (Fig 2f-g). AAV2_wt and in particular AAV9_wt targeted many cells outside the v-SVZ, especially in the medial and dorsal wall of the lateral ventricles, whereas the striatum was not targeted (Fig 2f,g and data not shown). In contrast to the wt capsids, AAV1_P5 and AAV9_A2 demonstrated a significantly higher tropism towards the v-SVZ (Fig 2i). AAV1_P5 showed the most unique tropism with 98% of all tdTomato-labeled cells lying along the v-SVZ. In addition, transduction rates of overall cells also differed between the four capsids. AAV1_P5 and AAV9_A2 exhibited the fastest kinetics and most robust rate of transduction, with AAV1_P5 transducing the largest number of cells at 5 dpi as compared to the other capsids (Fig 2j). The overall number of transduced NSCs became similar at 7 dpi for all capsids except AAV2_wt (Fig 2j). Nevertheless, AAV9_wt mostly targeted cells lying outside the ventricular wall that we clearly identified as neurons based on their morphology. By contrast, AAV1_P5 and AAV9_A2 exhibited a selective tropism for the v-SVZ mainly targeting NSCs/TAPs (Sox2^+^/GFAP^+/−^/S100b^−^) as well as ependymal cells (Sox^+^/S100b^+^; Fig 2h,j).

Along the wall of the v-SVZ, ependymal cells are organized in a so-called pinwheel architecture with NSCs in the center^53^. Within these structures, ependymal cells outnumber NSCs, explaining why AAV1_P5 and AAV9_A2 transduce more ependymal cells overall. A recent report using single cell transcriptomics and fate-mapping of ependymal cells demonstrates their inability to generate progeny even after growth factor administration or brain injury^54^. This ensures that progeny labeled with AAV1_P5 or AAV9_A2 stems from NSCs. However, manipulated ependymal cells communicate with neighboring NSCs and might indirectly change the progeny of these NSCs. To address this, strategies to de-target ependymal cells, such as using a miRNA-regulated viral vector^55,56^ or an NSC-specific promoter, might be of use. Taken together, our data demonstrate a unique tropism and fast targeting of NSCs/TAPs and ependymal cells within the v-SVZ by AAV1_P5 and AAV9_A2.

To select the best candidate between AAV1_P5 and AAV9_A2 regarding NSC transduction efficiency, we performed FACS analysis of the v-SVZ and OB of injected mice. 2 months old C57BL/6N mice were injected with 10^10^ vg in 10 μl of either AAV1_P5 or AAV9_A2 capsids containing the eYFP reporter under the CMV promoter, as these were the capsids used for the barcoded libraries. 6 days after injection, mice were sacrificed and NSCs with their progeny from the v-SVZ and the OB neuroblasts were analyzed by FACS quantification (Fig S3c and S6a,b). By determining the fraction of YFP^+^ cells among these cell types, we calculated the labeling efficiency of the different viruses. Our results show that AAV1_P5 has a higher labeling efficiency for NSC (11.19%) than the AAV9_A2 capsid (2.95%) (Fig S3d). This higher transduction efficiency could also be seen for qNSC, aNSC, TAPs and NBs from the SVZ (Fig S3d). This prompted us to proceed with the AAV1_P5 capsid for further experiments. Of note, the overall low number of detected YFP-positive cells is due to the lower sensitivity of FACS analysis for YFP-expressing cells as compared to mCherry or tdTomato, as previously shown (Tlx-YFP vs. tdTomato-YFP in^57^).

In order to test the ability of directly AAV1_P5 transduced NSCs to generate progeny, freshly isolated NSCs from tdTomato-flox mice were transduced with AAV1_P5 expressing Cre recombinase under the control of a CMV promoter (CMV_Cre). Thereafter, transduced cells were transplanted into the v-SVZ of C57BL/6N wt mice (Fig S4a). After 35 days, tdTomato-positive neurons were present in the GCL of the OB (Fig S4b-d). In summary, transduction of NSCs by AAV1_P5 *ex vivo* does not interfere with their capability to self-renew and differentiate into OB interneurons.

In order to fully characterize the identity of AAV1_P5 transduced cells, as well as potential changes arising by the AAV-transduction itself, we profiled these cells and untransduced ones from the same mouse by single cell RNA sequencing (scRNA-seq). To this end, three months-old eYFP-reporter mice (TiCY, Tlx-CreERT2-YFP mice^58^) were injected with 10^9^ vg/mouse AAV1_P5 harboring the CMV_Cre construct. Upon transduction, Cre recombinase causes the excision of a transcription terminator upstream of eYFP, which leads to eYFP expression. Transduction also causes excision of the neomycin resistance gene NeoR (Fig 3a, top). 37 days post injection, we isolated cells from the v-SVZ and other brain regions as schematically depicted in Fig 3a. More precisely, we isolated labeled cells of the v-SVZ and the striatum, rostral migratory stream (RMS) and OB, here referred to as rest of the brain (RoB). To capture the remaining unlabeled cells of the NSC-lineage in the v-SVZ, we also isolated GLAST^+^ v-SVZ cells. Two samples of two pooled mice each were subjected to scRNA-seq (Fig 3a and Fig S4e-g). Initial inspection of the resulting 4,572 single cell transcriptomes revealed a segregation of proliferating cells as indicated by the expression of the proliferation marker protein Ki-67 (Mki67) and canonical markers of G2/M and S phase (Fig S4h,i). After mitigating the effects of phase heterogeneity by regression, we obtained a continuous trajectory ranging from NSCs to late NBs / immature neurons (Fig 3b). Surprisingly, few eYFP^+^ off-target cells (sample #1: 9.7%; sample #2: 2.7%) were captured, consisting of mostly ependymal cells. Cells isolated from RoB are located at the very end of this trajectory (Fig S4j).

**Figure 3:**
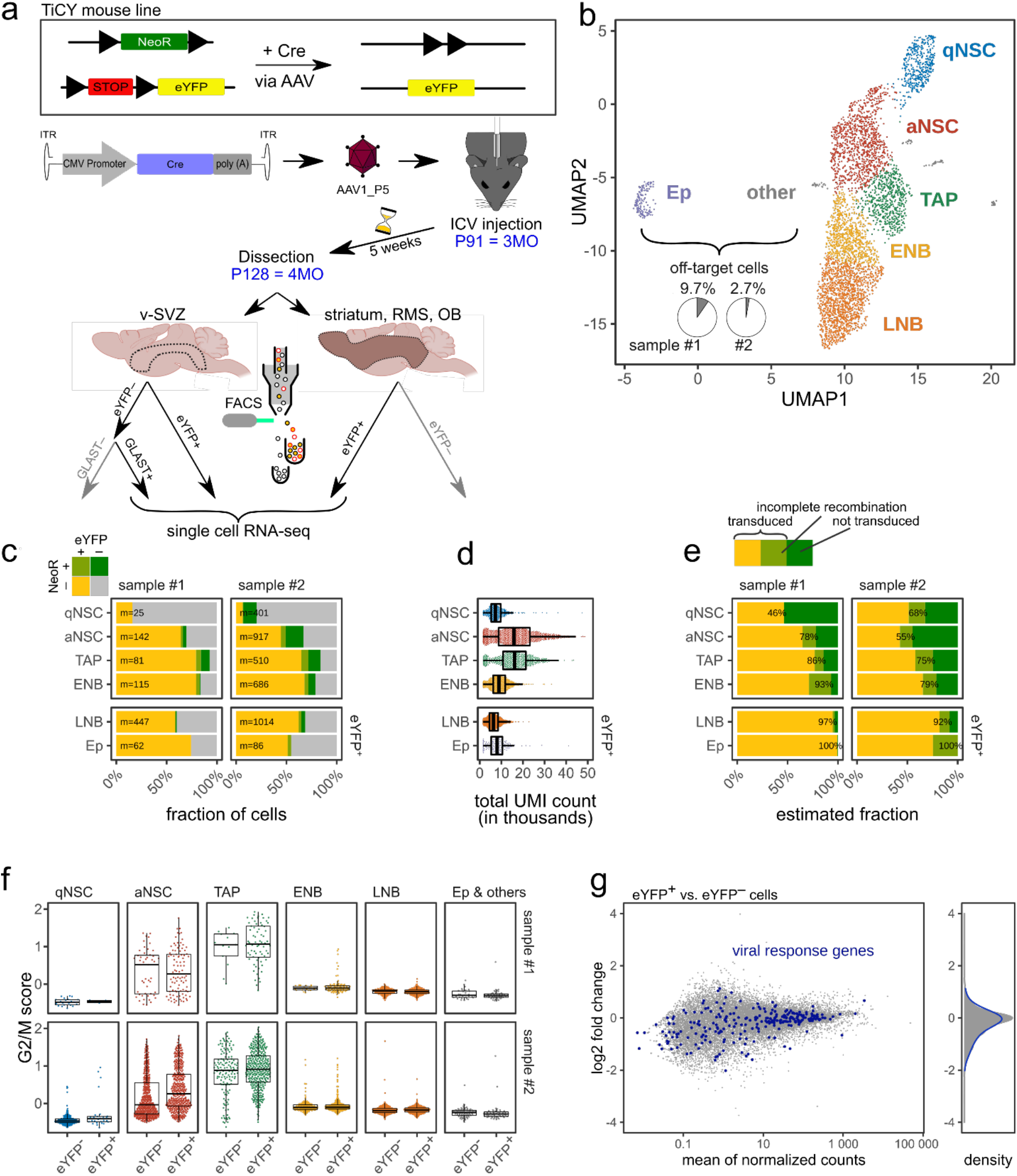
single cell RNA-seq reveals transduction of cells of the adult NSC lineage by AAV1_P5. **a** Experimental outline of labeling, isolation and single cell RNA sequencing (scRNA-seq) of the adult NSC lineage using the AAV1_P5 capsid. Top panel: Untransduced cells from the TiCY mouse line express NeoR. Cre-mediated recombination induces the expression of eYFP and the loss of NeoR expression. AAV1_P5 loaded with Cre was delivered to the lateral ventricle of P91 TiCY mice. After 5 weeks, all labeled (eYFP^+^) cells from the v-SVZ and the rest of the brain (striatum, rostral migratory stream [RMS] and olfactory bulb [OB]), as well as further unlabeled NSC lineage cells (GLAST^+^ from v-SVZ) were sorted and used for scRNA-seq. **b** 2D representation of the resulting 4,572 single-cell transcriptomes. Most cells form one continuous trajectory from qNSCs to early NBs (ENB; mostly from v-SVZ) and late NBs / immature neurons (LNB; mostly from rest of brain). Few off-target cells including ependymal cells (Ep) and others (gray) were captured. **c** Fraction of eYFP^+^ and NeoR^+^ single-cell transcriptomes by cell type (m: cells per group). **d** Total number of uniquely identified mRNA molecules (UMI count) per cell, separated by cell type. **e** Maximum likelihood estimate of the fraction of transduced cells, based on values in c and d. LNB and Ep were sorted by eYFP^+^ only and act as a control with an expected transduction rate of 100%. **f** Expression of G2/M phase marker genes across samples and cell types (clusters from b), distinguishing between eYFP^+^ and eYFP^-^ cells. **g** Left: MA plot of gene expression differences between eYFP^+^ and eYFP^-^ cells. Right: log2 fold change distribution for all genes (gray) and viral response genes (blue). Cre, Cre recombinase; ICV, intracerebroventricular; eYFP, enhanced yellow fluorescent protein; NeoR, neomycin resistance. (See also Figure S4)

Next, we sought to distinguish labeled (eYFP^+^ NeoR^-^) cells from unlabeled (eYFP^-^ NeoR^+^) cells in our single cell transcriptomes (Fig 3c, Fig S4k). As expected (Fig 3a, top), eYFP-expressing cells mostly do not express NeoR, and, *vice versa,* cells expressing NeoR mostly do not express eYFP. Only very few cells express both eYFP and NeoR (samples #1 and #2: 1.4% and 3.7%), possibly due to incomplete Cre-mediated excision. Transcripts of the viral Cre-recombinase, however, were rarely detected and mostly in early stages of the lineage but notably, also in very few cells at the end of the lineage, indicating an overall very low expression that prevents estimation of the dilution of viral transcripts along the lineage (Fig S4l). The floxed genes, eYFP and NeoR exhibited higher expression than the Cre transcript. eYFP was more readily detected than NeoR, but ultimately both genes suffered from the usual “dropout” in scRNA-seq, i.e. the failure to capture and/or detect transcripts^59^. For a substantial fraction of cells, neither NeoR nor eYFP was detected. The fraction of such undistinguishable cells was larger in cells with fewer total detected transcripts such as qNSCs and LNBs (Fig 3c and Fig 3d). To overcome this issue and estimate AAV1_P5 transduction efficiency while accounting for total transcript count per cell and the likely different expression strengths of eYFP and NeoR, we employed maximum likelihood estimation (Fig 3e, Methods). LNBs (mostly from eYFP^+^-sorted RoB) and ependymal cells (GLAST^-^) were used as controls since we know that almost all of these cells are transduced. Overall, we estimated a high transduction efficiency ranging from 46% to 93% for the cell types of the v-SVZ lineage and estimated 92% to 100% transduction in cells used as controls. Lastly, we assessed whether the transduced cells show transcriptomic differences arising from the viral transduction itself. Both eYFP^-^ and eYFP^+^ aNSCs and TAPs showed high expression of commonly used G2/M phase marker genes (Fig 3f), which suggests that transduction with AAV1_P5 does not affect proliferation. Differential gene expression analysis between eYFP^+^ cells and eYFP^-^ cells (Fig 3g) identified only 18 differentially expressed genes, indicating that AAV1_P5 transduction affects their transcriptome only mildly. Furthermore, we did not find any concerted upregulation of viral response genes in this comparison, or when comparing eYFP^+^ cells to eYFP^-^ NeoR^+^ cells (Fig S4m) or naive v-SVZ lineage cells from^2^ (Fig S4n). In conclusion, we have combined scRNA-seq with lineage tracing using AAV1_P5 and found that transduction does not affect the expression of proliferation markers and overall only minimally affects the transcriptomic readout.

We next tested whether the transduction efficiency could be further optimized by the selection of promoter and number of injected vg per mouse. To this end, we now packaged the CMV_Cre construct into the AAV1_P5 capsid and injected either 10^9^ vg per mouse as in Fig 2e-j, or an increased concentration of 10^10^ vg per mouse into tdTomato-flox mouse brains (Fig S5a). In all conditions, tdTomato-labeled cells were detected at high numbers in the v-SVZ, confirming specific v-SVZ targeting by the AAV1_P5 capsid (Fig S5b-d). Transduction of cells was over 60 times higher with the CMV_Cre construct (319.9 cells per section, Fig S5d) than with CAG_Cre (4.8 cells per section, Fig 2j) when injecting 10^9^ vg per mouse. By increasing the number of injected vg from 10^9^ to 10^10^, we were able to further increase the number of labeled cells (Fig S5d) including NSCs / TAPs and ependymal cells (Fig S5f-g). However, the increased viral load also moderately increased the proportion of labeled cells located outside of the v-SVZ (Fig S5e). Overall, we found that increased viral load resulted in higher labeling efficiency as expected, but at the cost of some regional specificity. This trade-off must be considered when designing future experiments, e.g. when targeting cells outside the v-SVZ must be absolutely avoided it is advisable to inject a lower amount of vg. Furthermore, the CMV promoter greatly outperformed the CAG promoter in our experiment. This result differs from previous studies overexpressing plasmids via *in utero* electroporation in the mouse brain, which showed a higher efficiency of the CAG than the CMV promoter^60,61^. We conclude that the CMV promoter should be preferred over CAG when using AAV1_P5, injecting 10^10^ vg per mouse or alternatively 10^9^ when regional specificity is crucial.

We finally assessed the neurogenic function of transduced NSCs *in vivo.* To this end, we assessed the number of transduced NSCs in the v-SVZ and their neuronal progeny in the OB. 10^10^ vg/mouse of AAV1_P5 harboring the CMV_Cre construct were injected into the lateral ventricles of tdTomato-flox mice and at 35 dpi, the number of labeled NSCs in the v-SVZ and OB interneurons was assessed (Fig 4a). We observed a high heterogeneity in the number of labeled cells probably due to differences in the injection site. One set of animals exhibited a lower number of labeled cells in the SVZ and OB than the other (Fig 4b). While a trend towards a reduced number of NSCs/TAPs at 35 dpi was detectable, NSCs still remained in the v-SVZ at this late time point (Fig 4c), suggesting that AAV1_P5 also targeted qNSCs. To estimate the extent of targeting of the NSC compartment, we took advantage of our previously developed mathematical modeling framework for stem cell dynamics of v-SVZ^2^. First, we extended our previously established model and calibrated it to the experimentally observed dynamics of TAPs and OB neurons (see Supplementary Material: Mathematical Modeling). Instead of fitting the model to average cell counts across mice, we subdivided the data into two groups, with higher and lower labeling, as animals with high labeling in the v-SVZ exhibited a much higher number of labeled cells in the OB than animals with lower labeling (Fig 4d-e). Fitting of the model to the data, assuming that viral transduction does not affect cell kinetics and that the observed heterogeneity comes from different numbers of initially labeled NSCs and TAPs, the model indicates that approximately 57% of NSCs are labeled in the high-label group and 26% of NSCs in the other group (see Supplemental Material). Moreover, the model indicates that in the low-labeled group, barely any TAP would be labeled at initial time, whereas in the other group a higher number of TAPs is initially labeled. Finally, we employed our model to address whether the observed labeling would arise from direct targeting of qNSCs, aNSCs or both. To this end, we simulated two scenarios where either only qNSCs or only aNSCs are targeted (Fig 4f). Our simulation indicates that the ratio of labeled qNSCs to aNSCs reaches the same value in both scenarios after approximately four days, due to transitions between the quiescent and active state. Altogether comparison of model fit to data is in line with the hypothesis that the number of initially transduced NSCs and TAPs differs between the two groups, that the cell dynamics exhibited by transduced cells are comparable to non-transduced cells and that AAV5_P1 can target up to 57% of the NSC pool.

**Figure 4:**
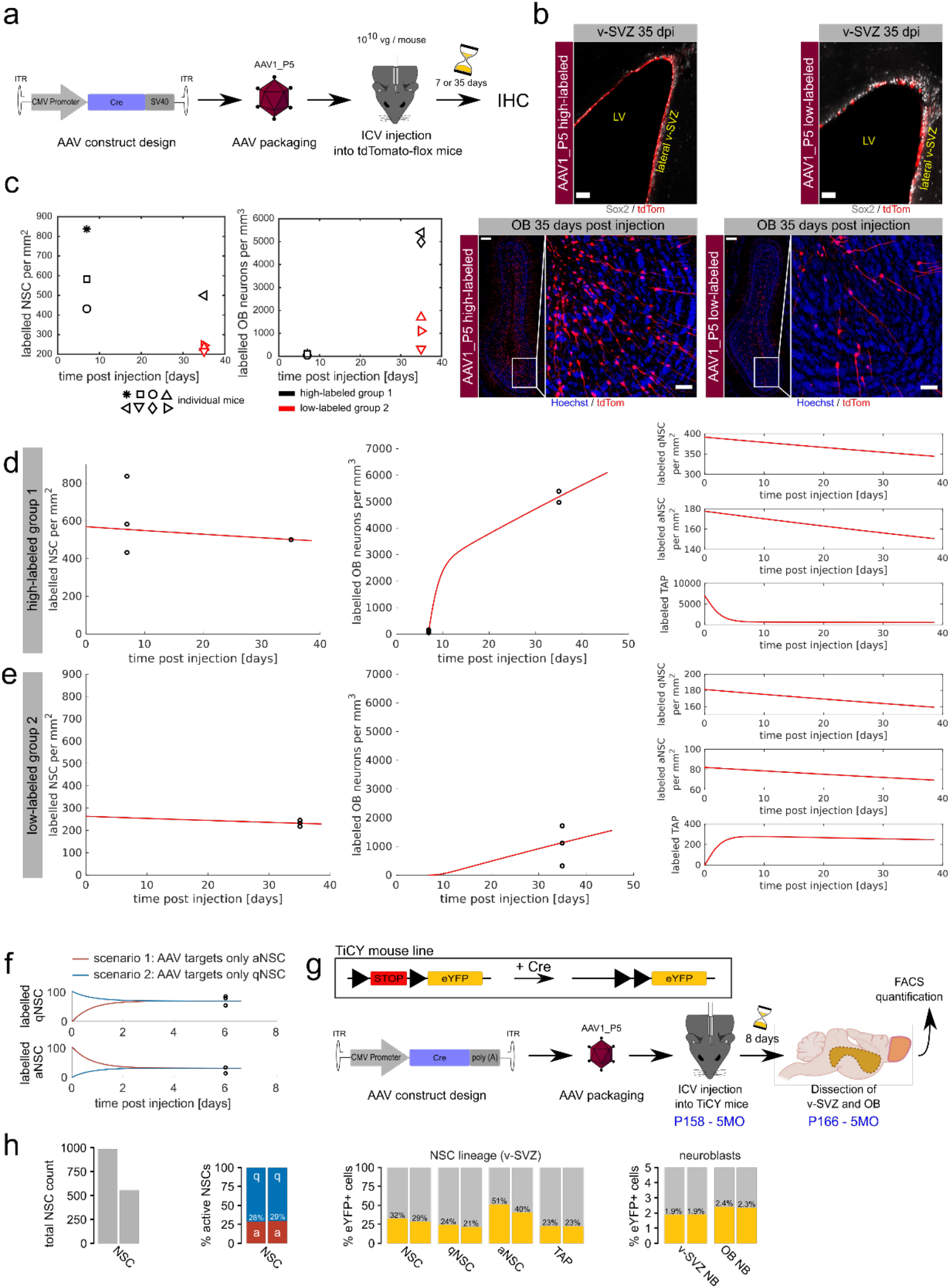
AAV1_P5 targets qNSCs and the choice of promoter and viral load determines the number of generated OB neurons. **a** Schematic illustration of the experimental outline to test v-SVZ labeling with at different time points. **b** IHC of the v-SVZ (scale bar 50μm) and OB (scale bar 200μm and 50μm) in the high-labeled and low labeled group 35 days post AAV injection. **c** Time dynamics of labeled cells. Each mouse is identified by one symbol. Due to the heterogeneity among individual mice, each mouse was assigned to one of two groups. The color of the symbols indicates to which group the respective mouse belongs. **d,e** Comparison of model fit and data. **d** compares the fit to data from the high-labeled group 1 and **e** compares to data from low-labeled group 2. The model was fit to both groups simultaneously. Only the number of initially labeled NSCs and TAPs differs between the two groups. **f** Redistribution of labeled NSC between the active and the quiescent state. We compare two scenarios. In the first scenario (red lines) the virus targets only aNSC. In the second scenario (blue lines) the virus targets only qNSCs. After 4 days the number of labeled aNSC are identical for both scenarios (lower panel). The same applies to the number of labeled qNSCs (upper panel), since aNSC can become quiescent after division and qNSCs can become activated. Black dots indicate FACS quantifications of NSCs labeled by the AAV1_P5_YFP adenovirus (as shown in Fig S3c,d). Virus injection took place at time 0. **g** Experimental layout of FACS quantification of TiCY mice to analyze labeling efficiency of the v-SVZ and olfactory bulb using AAV1_P5_Cre. **h** Quantification of FACS events: Total NSC count in the v-SVZ; proportion of active to quiescent NSCs; proportion of eYFP^+^ NSCs and TAPs; and proportion of eYFP^+^ neuroblasts in the v-SVZ and olfactory bulb. Cre, Cre recombinase; SV40, Simian-Virus 40 polyA signal; ICV, intracerebroventricular. (See also Figure S4 and S5).

To validate the model prediction of the label efficiency of the AAV1_P5 vector, we performed a FACS quantification experiment to directly assess the percentage of NSC and progeny that is labeled by the virus 8 days after injection (Fig 4g). 5 months old TiCY mice were injected with 10^9^ vg/mouse of AAV1_P5 harboring the CMV_Cre construct. FACS quantification analysis was performed as described previously (Fig S3c-d, S6a,b) and the results show 30.46% labeling efficiency for NSCs (Fig 4h, mean eYFP^+^-percentage of both samples), which is close to the 26% labeling efficiency predicted by the mathematical model (Supplementary Material: Mathematical Modeling, Section 3.2). The model also showed a good fit when applied to the FACS quantification experiment performed to choose the best candidate between AAV1_P5 and AAV9_A2. Moreover, the prediction of a high labeling group was validated by the observed labeling rate in the single cell transcriptomics analysis (see Supplementary Material: Mathematical Modeling).

## Discussion

Altogether, in this study we have performed barcode-based *in vitro* and *in vivo* high-throughput screenings of two libraries of wt and engineered AAV capsids^50^. Targeting of NSCs and especially qNSCs has only been demonstrated in the hippocampal dentate gyrus with the capsid AAV r3.45^62^ and the African green monkey isolate AAV4^63^, as well as recently in the v-SVZ using the newly engineered AAV variant SCH9^39^.

Here, we have identified two lead candidates for efficient targeting of NSCs *ex* and *in vivo.* We particularly characterized the novel capsid AAV1_P5 as highly region-specific at targeting cells of the v-SVZ layer, including ependymal cells and NSCs, by IHC, FACS quantification and scRNA-seq. We moreover show by IHC and scRNA-seq that NSCs targeted with AAV1_P5 were not noticeably affected in their migration and transcriptome and readily generated OB neurons. Furthermore, we demonstrate that the engineered capsid AAV1_P5 also labels qNSCs. We propose that qNSC labeling can not only be achieved by direct targeting of qNSCs, but also indirectly through transduction of aNSCs that would later give rise to qNSCs. Indeed, based on mathematical modeling of FACS counts, we predict that labeled cells redistribute between those states within less than one week. Therefore, the initial labeling proportion of quiescent to active NSCs is not crucial when stem cell dynamics are observed on a longer time scale.

AAV1_P5 clearly shows a tropism for the v-SVZ and is unable to migrate further away from this region, but the molecular mechanism for this tropism is unknown. It was previously shown that the SCH9 variant binds heparan sulfate proteoglycans and galactose, both of which are present on NSCs in the v-SVZ^39^. To date, there are only few other cases where such mechanisms underlying altered viral properties of synthetic AAV capsids have been successfully elucidated^64–67^. One example is the use of αvβ8 integrin as receptor for a keratinocyte-specific AAV2^64^. Another example was reported by several labs who have recently identified an interaction of AAV-PHP.B (a peptide-modified AAV9) with the GPI-linked protein LY6A^65–67^. Other than these, however, the receptors or interactions that are targeted by peptide-engineered or shuffled AAV variants typically remain enigmatic, as do the intracellular mechanisms underlying their novel features. Hence, identifying the receptor for AAV1_P5 will be the subject of future studies. In this looming work, it will then also be interesting to study whether AAV1_P5 interacts with other host cell factors which have been identified over the years as critical for transduction with wild-type capsids, such as the widely used AAV receptor AAVR^68^ or intracellular elements such as the proteasome^69^.

As a proof-of-concept, we show that AAV1_P5-labeling can be combined with scRNA-seq to characterize the transcriptomes of NSCs and their progeny from different brain regions. This paves the way for more complex lineage tracing experiments *in vivo.* Recent studies have used CRISPR-Cas9-induced genomic scars combined with scRNA-seq to enable clonal lineage tracing in embryonic development^70,71^. AAVs could be used to induce genomic scars in specific cells at specific time points to enable clonal lineage tracing in adult tissues. We use our scRNA-seq data to further corroborate our assessment that NSCs are efficiently targeted and remain functional after transduction. Future studies using electrophysiology are required to assess whether the progeny generated by transduced NSCs is fully functional and able to integrate into the neuronal circuits of the OB. We have identified the combination of CMV promoter and AAV1_P5 capsid as ideally suited to efficiently transduce NSCs in the v-SVZ. In addition, we think that future experiments will be needed to unravel and understand the mechanisms governing the properties of our candidates. Altogether, we believe that our study opens tantalizing avenues to genetically modify NSCs in their *in vivo* environment for the treatment of CNS disorders or brain tumors.

## Material and Methods

### Animals

In this work, the mouse lines C57BL/6N, TdTom-flox [B6-Gt(ROSA)26Sortm14(CAG-tdTomato)Hze] and TiCY [B6-Tg(Nr2e1-Cre/ERT2)1Gsc Gt(ROSA)26Sortm1(EYFP)CosFastm1Cgn/Amv] were used. All mice were male and were age-matched to eight weeks, except for TiCY mice, which were five months old (for FACS quantification) and three months old (for scRNA-seq). Animals were housed in the animal facilities of the German Cancer Research Center (DKFZ) at a 12 h dark/light cycle with free access to food and water. All animal experiments were performed in accordance with the institutional guidelines of the DKFZ and were approved by the “Regierungspräsidium Karlsruhe”, Germany.

### AAV Vector Production

The production of the AAV barcoded library was done as previously published^72,73^ with some modifications: 159 distinct barcodes were inserted into the 3’ untranslated region of a yellow fluorescent protein (YFP) reporter under the control of a cytomegalovirus (CMV) promoter and encoded in a self-complementary AAV genome. Each of the barcodes was assigned to one AAV capsid from a total of 183 variants, which are described in more detail in the accompanying manuscript by ^50^. Altogether, this library production included 12 AAV-wild types (AAV1 to AAV9, AVVrh.10, AAVpo.1, AAV12) and 94 peptide display mutants, 71 capsid-chimeras, which were created by DNA family shuffling. Isolation of synthetic capsids were performed in specific tissues or in our recent screens of AAV libraries in cultured cells, in mouse liver tissue or muscle^74^. These synthetic capsids include a set of 12 AAV serotypes, that were previously modified by insertion of over 20 different peptides in exposed capsid loops and that were recently characterized in established or primary cells^74^. In the work of ^50^, all barcoded capsids were pooled in different combinations to finally obtain three distinct libraries (#1, #2 (not used in the present work) and #3), with 91, 82 and 157 variants. Further details on library composition are found in the Supplement of ^50^. All capsid variants are detailed in Table S4. HEK293T cells were cultured in DMEM (Gibco) supplemented with 10% fetal bovine serum (Merck), 1% penicillin/streptomycin (Gibco, 10000 U/ml pen, 10000 μg/ml strep) and 1% L-Glutamine (Gibco, 200 mM) at 37°C and 5% CO2. AAV vectors were produced by seeding HEK293T cells (4.5×10^6^ cells per dish) on 90-150 Ø15cm tissue culture dishes (Sigma). Two days later, we performed a polyethylenimine (PEI; Polyscience) triple transfection by mixing 44.1 μg (3×14.7 μg) DNA of i) a plasmid containing the recombinant AAV genome of interest ii) an AAV helper plasmid carrying AAV *rep* and *cap* genes and iii) a plasmid providing adenoviral helper functions for AAV production in a total volume of 790 μl H_2_O per culture dish. Separately, PEI (113.7 μg) and H_2_O were mixed in a total volume of 790 μl per dish and NaCl (300 nM) was added 1:1 to both, PEI or DNA solution. PEI was added dropwise to DNA and incubated for 10 min at room temperature, before finally adding the DNA/PEI mixture to the culture dish. Three days later, cells were scraped off in the media and collected by centrifugation (400 g, 15 min). The pellet was dissolved in 0.5 ml virus lysis solution (50 mM TrisHCl; Sigma), 2 mM MgCl2 (Sigma), 150 mM NaCl (ThermoFisher; pH 8.5) and was immediately frozen at −80°C. In total, 5x freeze-thaw cycles were performed with the cell pellet prior to sonication for 1 min 20 sec. The cell lysate was treated with Benzonase (75 U/μl; Merck) for 1h at 37°C, followed by a centrifugation step at 4000 g for 15 min. CaCl2 was added to a final concentration of 25 mM and the solution was incubated for 1 h on ice, followed by centrifugation at 10000 g for 15 min at 4°C. The supernatant was harvested and a ¼ volume of a 40% polyethylene glycol (PEG 8000; BioChemica) and 1.915 M NaCl (ThermoFisher) solution was added prior to incubation for 3 h on ice. After centrifugation for 30 min at 2500 g and 4°C, the pellet was dissolved in resuspension buffer (50 mM HEPES; Gibco), 0.15 M NaCl (ThermoFisher), 25 mM EDTA (Sigma) and was dissolved overnight. The solution was then centrifuged for 30 min at 2500 g and 4°C, and the supernatant was mixed with cesium chloride (CsCl; Sigma) to a final concentration of 0.55 g/ml. The refractive index was adjusted to 1.3710 using additional CsCl or buffer, as needed. Next, the vector particles were purified using CsCl gradient density centrifugation. Fractions with a refractive index of 1.3711 to 1.3766 comprising DNA-containing AAV particles were pooled and dialyzed against 1x PBS with a Slide-A-Lyzer dialysis cassette according to the manufacturer’s instructions (ThermoFisher). Subsequently, the samples were concentrated by using an Amicon® Ultra Centrifugal Filter (Millipore; 100000 NMWL) following the manufacturer’s instructions. The volume of the samples was reduced to 250-300 μl. AAV vectors were finally aliquoted and stored at −80°C.

The production of the AAV1_P5_YFP and AAV9_A2_YFP viruses for the FACS analysis experiment was done as described above, with the only modification that the vectors were purified using two Iodixanol gradients. Of note, the barcoded AAV library construct as well as the YFP-construct were engineered as double-stranded AAV vectors. The constructs for CAG_Cre::GFP and CMV_Cre were engineered as a single-stranded AAV vector.

### AAV Vector Titration

AAV vectors were titrated using quantitative real-time PCR (qRT-PCR) as described in^75^. For the CAG_Cre::GFP construct, the primers and probe GFP_fwd, GFP_rev and GFP_probe were used, while Cre_fwd, Cre_rev and Cre_probe were used for the CMV_Cre construct (Table S1). The qPCR was performed on a C1000 Touch Thermal Cycler equipped with a CFX384 Real-Time System (Bio Rad) with the following conditions: initial melting for 10 min at 95°C, followed by 40 cycles of denaturation for 10 s at 95°C and annealing/extension for 30 s at 55°C. A standard curve was considered as reliable when R^2^ was greater than 0.985.

### Stereotactic Injection

AAV vectors were stereotactically injected into the lateral ventricle by using the following coordinates calculated to bregma: Anterior-posterior (AP) −0.5 mm, Medio-lateral (ML) −1.1 mm, Dorso-ventral (DV) 2.4 mm. Mice received either 10^9^ or 10^10^ vg/mouse in a total volume of 10 μl. The AAV libraries were stereotactically injected into the lateral ventricle by using the following coordinates calculated to bregma: AP −0.5 mm, ML −1.1 mm, DV 2.4 mm. Mice received 4 x 10^10^ vg/mouse in a total volume of 2 μl. *Ex vivo* manipulated cells (7000 FACS events) were injected into two areas of the v-SVZ using the following coordinates calculated to bregma: AP 0.7 mm, ML 1.6 mm, DV 2 mm and AP 0 mm, ML 1.7 mm, DV 2 mm.

### Cell Isolation and *in vitro* Cultivation

The lateral v-SVZ was micro-dissected as whole mount as previously described^76^. Tissue of single mice was digested with trypsin and DNase according to the guidelines of the Neural Tissue Dissociation Kit (T) (Miltenyi Biotec) using a Gentle MACS Dissociator (Miltenyi Biotec). Cells were cultured and expanded for 8-12 days in Neurobasal medium (Gibco) supplemented with B27 (Gibco), heparin (Sigma), glutamine (Gibco), Pen/Strep (Gibco), EGF (PromoKine) and FGF (PeloBiotech) as reported in ^77^.

### *In vitro* Transduction of Cultured NSCs

For RNA sequencing, NSCs were seeded in 48-well plates (Greiner Bio-One) and incubated overnight. AAV library #1 or library #3 (same libraries as in ^50^, multiplicity of infection (MOI): 10000) were added to the media and remained for the duration of seven days. For IHC, Labtek chambers (ThermoFisher) were coated with PDL (Sigma) / Laminin (Sigma) and NSCs were seeded at a density of 2×10^4^ cells per cm^2^ overnight. AAVs were added (MOI: 10000) and remained in the media for 1, 3, 5 or 7 days.

### Single-cell transcriptomic profiling by 10X Chromium 3’ sequencing

### Stereotactic injection, single cell suspension preparation and sorting

Three months old TiCY mice were stereotactically injected into the lateral ventricle with 10^9^ vg of the AAV1_P5_Cre capsid. After 5 weeks of chase time, the mice were sacrificed and the SVZ, striatum, rostral migratory stream and olfactory bulb was isolated. The latter three tissues were pooled as a single tube and were named Rest of the Brain (RoB). From these tissues a single cell suspension was prepared as described before (Cell Isolation and in vitro Cultivation section). From the SVZ the cells sorted were eYFP^+^ (O4/CD45/Ter119 negative, eYFP positive) and, from the eYFP negative cells, only Glast^+^ cells. From the RoB only eYFP^+^ cells were sorted. The total number of sorted events for the 2 days of the experiment were 12000 for SVZ cells and 5800 for cells of the RoB. 2 TiCY mice were pooled for each sorting day. All the cells were sorted in a volume of 50 μl of Fetal Calf Serum (FSC) 10% in PBS, from which 45 μl were used for loading the Chromium Next GEM Chip G.

### Library preparation, sequencing, and mapping

One library per each sorting day was prepared by following the manufacturer’s protocol (Chromium Next GEM Single Cell 3’ v3.1) and sequenced on a NovaSeq 6K PE 100 S1.

In order to quantify eYFP and NeoR (Neomycin / Kanamycin resistance gene) expression, entries for these transgenes were manually added to the FASTA and GTF files of the mouse reference genome mm10-3.0.0 provided by 10X Genomics. scRNA-seq reads were pseudoaligned and further processed with kallisto|bustools^78,79^ to generate a gene×barcode count matrix.

### Computational analysis of single cell RNA-seq data

Cell barcodes with less than 1500 UMIs or more than 15% mitochondrial reads were filtered and the remaining cells were further analyzed in Scanpy v1.5.1^80^. We used Scanpy to calculate G2/M and S phase scores for all cells, based on their expression of G2/M and S phase marker genes from ^81^. These scores were then regressed out of the count data, to reduce the influence of the cell cycle on clustering. The first 50 principal components of 3324 highly variable genes were used for 2D visualization with UMAP (n_neighbors=35) and cell clustering with the Leiden algorithm (resolution=0.5). Cell clusters were assigned to cell types based on the expression of marker genes as previously described in^2^. To identify the location of cells from RoB, kernel density estimates of cell density in 2D UMAP space were calculated for both samples. Since sample #1 contains more RoB cells and sample #2 contains more v-SVZ cells, we subtracted both densities to highlight cells that most likely stem from RoB (orange cells in Fig S4h).

In order to estimate transduction efficiency from scRNA-seq data, we use the following model, based on the usual approach of modeling RNA-seq counts by the negative binomial (NB) distribution:

For non-transduced cells, we assume that they express NeoR such that an expected fraction *μ*_R_ of all their mRNA transcripts originate from this gene. For each individual cell *j*, the actual expression strength 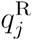 of the gene varies around this expectation according to a gamma distribution with mean *μ*_R_ and variance *α*_R_/*μ*_R_. The observed number of UMIs is then modelled as a Poisson variable: 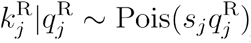 where *s_j_* is the total UMI count for cell *j,* summed over all genes. Marginalizing out 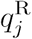, we 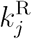 find to follow a NB distribution with mean *s_j_μ*_R_ and dispersion *α*_R_. As we are looking at a non-transduced cell, the UMI count 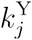 for eYFP is, of course, zero.

Similarly, we write 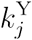, *μ*_R_ and *α*_Y_ for the corresponding quantities of eYFP, expressed by transduced cells. For a fully transduced cell *j,* we therefore have 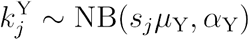, but 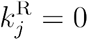. For transduced cells with incomplete or heterozygous Cre-mediated excision, we should see both genes expressed, but will model the expression strength to be only half as strong.

The likelihood of observing UMI counts 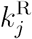 and 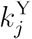 for a given cell *j* therefore depends on the parameters just mentioned as well as on the probabilities *Pν* that the cell is not transduced, *p*_T_ that it is fully transduced, and *p*_P_ = 1 – *p*_U_ – *p*_r_ that it is partially transduced. We write the likelihood as

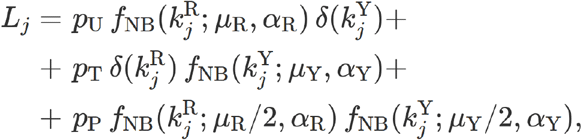

where *f*_NB_ (*k*; *μ, α*) is the probability to observe *k* counts under a negative binomial distribution with mean *μ* and dispersion *α*, and *δ* is the zero indicator function, i.e., *δ*(*k*) = 0 for *k* ≠ 0 but *δ*(0) = 1.

Given all the *k_j_* and *s_j_*, we obtain estimates for the transduction efficiency *p*_T_ and for *p*_U_ and *p_P_* as well as for the nuisance parameters *μ_R_*, *α_R_*, *μ_Y_*, and *α_Y_* by numerically maximizing the log likelihood 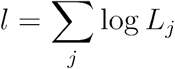 using the R function optim.

We mention two technical details: First, in order to give all optimization parameters full domain over all of ℝ, we used parameter transformations in the optimization, namely exponentiating the *μ*_S_ and *α*s. and logit-transforming the probabilities *p* and *q* obtained from reparametrizing *p_T_ = *p*(1 – *q*), *p_U_** = 1 – *p*, *p_P_* = *pq*. Second, in order to improve identifiability in case of low values for *p_U_*, we enforced a minimum value for *μ_R_* by adding to the likelihood a penalty term *f*_pty_ (*μ_R_*), where *f*_pty_ = 1/(1 + *e*^9×10^5^*x*–9^)is a sigmoid that vanishes for *μ_R_* ≳ *μ*_*R*min_ = 2×10^−5^.

Differential gene expression was assessed by summing UMI counts of cells within a group to yield pseudobulk samples for testing in DESeq2 v1.29.7^82^. eYFP^+^ cells were tested against both eYFP^-^ cells and eYFP^-^ NeoR^+^ cells. Testing eYFP^+^ vs. eYFP^-^ has the advantage of greater statistical power due to higher cell numbers, but some eYFP^-^ cells may be transduced cells with eYFP-dropout. Thus, we performed both comparisons, yielding similar results. To account for the unequal distribution of eYFP^+^ and eYFP^-^ cells along the lineage (Fig S4h), pseudobulk groups were formed per cluster and sample, and the cluster identity was added as a covariate in DESeq2. To enable comparison of v-SVZ cells from ^2^ with our eYFP^+^ cells, both datasets were integrated with Seurat’s SCTransform integration workflow^83^ using our cells as reference. The integrated dataset was clustered and differential expression was assessed as above, using the shared clusters as covariate. Genes with the gene ontology term “GO:0009615 - response to virus” were highlighted.

### FACS-Sorting

Generation of single-cell suspension was performed as described in ^7^. Cells were stained with the following antibodies: O4-APC and O4-APC-Vio770 (Miltenyi; diluted 1:50), Ter119-APC-Cy7 (Biologend; 1:100), CD45-APC-Cy7 (BD; 1:200), GLAST (ACSA-1)-PE (Miltenyi: 1:20), CD9-eFluor450 (eBioscience: 1:300), Alexa647::EGF (Life Technologies, 1:100), PSA-NCAM-PE-Vio770 (Miltenyi; 1:75), Prominin1-PerCP-eFluor 710 (eBioscience; 1:75), CD24-PE-Cy7 (eBioscience; 1:75), and Sytox Blue (Life Technologies, 1:1000). For RNA sequencing, cells were directly sorted into 100 μl of the PicoPure RNA Isolation Kit (ThermoFisher) extraction buffer. For *ex vivo* transduction, NSCs were sorted into growth-factor-free NBM medium.

### FACS Analysis of AAV-injected mice

FACS Analysis for testing the transduction efficiency of the candidate viruses was performed by two methods. The first method consisted of injecting 5 months old TiCY mice with the AAV1_P5_Cre virus and after 8 days SVZ and OB cells were FACS sorted and analyzed (Fig 4g-h). In the second method we injected 2 months old C57BL/6N mice with AAV1_P5_YFP and AAV9_A2_YFP viruses and analyzed them after 6 days (Fig S3c-d).

For FACS Quantification of AAV-injected NSC/Progeny, cells were sorted with the following antibodies: O4-APC-Vio770 (Miltenyi; diluted 1:100), CD45-APC-Cy7 (BD; 1:200), Ter119-APC-Cy7 (Biologend; 1:100), GLAST (ACSA-1)-PE (Miltenyi: 1:50), Prominin1-APC (eBioscience, 1:75), PSA-NCAM-PE-Vio770 (Miltenyi; 1:50), Texas-Red::EGF (Life Technologies, 1:75).

### *Ex vivo* Treatment of NSCs

FACS-sorted NSCs were transduced with AAV (MOI: 10000) and incubated on ice for 2-3 h. Cells were centrifuged for 15 min at 300 g, 4°C and were washed twice with PBS. The pellet was dissolved in 4 μl PBS.

### RNA Isolation and cDNA Synthesis

RNA was isolated by using the PicoPure RNA Isolation Kit (ThermoFisher). For RNA isolation of *in vitro* transduced cells, 1500 cultured NSCs per set were lysed in 100 μl extraction buffer. For isolation of FACS-sorted *in vivo* transduced cells, batches of 500 cells or less were generated and were lysed in 100 μl extraction buffer. Up to 6 batches (2500 cells) were obtained per set, depending on the cell type (Table S2, Table S3). The cell-containing extraction buffer was incubated for 30 min at 42°C and the lysate was frozen at −80°C to increase the amount of isolated RNA. The cell lysate was mixed 1:1 with 70% ethanol and RNA was extracted according to the guidelines of the PicoPure RNA Isolation Kit (ThermoFisher). RNA was dissolved in 11 μl nuclease-free H_2_O. The cDNA synthesis was performed as described in^84^ by using Locked Nucleic Acid-TSO (Table S1) and by using either 14 cycles for *in vitro* cultured NSCs, 15 cycles (>300 cells per batch) or 16 cycles (<300 cells per batch) for FACS-sorted *in vivo* transduced cells for the cDNA enrichment step. After purification^84^ using AMPure XP beads (Beckman Coulter), cDNA was dissolved in 10 μl H_2_O.

### Barcode Amplification PCR and NGS Library Preparation

Barcodes were PCR-amplified by using 10 ng cDNA as input material. Therefore, the PCR primers Bar_fwd and Bar_rev that bind up and downstream of the 15bp long Barcodes within the according cDNA were engineered and the Phusion High-Fidelity DNA Polymerase (ThermoFisher) was used according to its manual in combination with 10 mM dNTPs (ThermoFisher) (Table S1). The PCR was performed on a T100 Thermal Cycler (Bio Rad) with the following conditions: initiation for 30 s at 98°C, followed by 35 cycles of denaturation for 10 s at 98°C, annealing/extension for 20 s at 72°C and a final step for 5 min at 72°C. The result was a 113bp long PCR amplicon that includes the Barcode with its 15bp long random DNA-sequence. The PCR amplicon was AMPure XP bead-purified (Beckman coulter)^84^ with a bead:sample ratio of 0.8:1 in the first round and 1:1 in the second round. After this step, the samples were enriched for the Barcode containing amplicon and of course the samples potentially contained the range of up to 157 different AAV Barcodes which were initially used. Next, 10 ng or 15 ng (library #1 or #3, respectively) of PCR amplicon was used for NGS library preparation with the NEBNext ChIP-Seq Library Prep Reagent Set for Illumina (NEB) for samples from library #1 and the NEBNext Ultra II DNA Library Prep Kit for Illumina (NEB) for samples from library #3. Multiplexed libraries were generated by following the manual and by using the NEBNext Multiplex Oligos for Illumina (NEB). All multiplexed samples for library #1 and library #3 are listed in Table S2 and Table S3. For sequencing, up to 50% of PhiX were spiked in to increase the complexity of the library.

### Immunocytochemistry

Cells were washed 3x 5 min in PBS at room temperature, followed by a 30 min blocking step in PBS^++^ (PBS with 0.3% horse serum (Millipore) and 0.3% Triton-X100 (Sigma)) at room temperature. Subsequently, the cells were incubated overnight in PBS^++^ containing primary antibodies at 4°C. Cells were washed in PBS for 3x 5 min at room temperature and were incubated with secondary antibodies in PBS^++^ for 1 h in the dark at room temperature. Afterwards, cells were washed 3x 5 min in PBS and were mounted with Fluoromount G (eBioscience). The following antibodies were used: chicken anti-GFP (Aves; 1:1000) and goat anti-mCherry (SICGEN; 1:1000). Nuclei were counterstained with Hoechst 33342 (Biotrend; 1:3000).

### Tissue Preparation

Animals were sacrificed by using an overdose of Ketamine (120 mg/kg) / Xylazine (20 mg/kg) and were subsequently transcardially perfused with ice-cold 20ml 1xHBSS (Gibco) and 10ml of 4% paraformaldehyde (Carl Roth). The brains were dissected and postfixed in 4% paraformaldehyde overnight at 4 °C. A Leica VT1200 Vibratome was used to cut the tissue in 50 μm (v-SVZ) or 70 μm (OB) thick coronal sections. From each mouse, three to six identical brain sections every 100 μm (v-SVZ) or 140 μm (OB) along the coronal axis were used for staining. Brain sections for staining the v-SVZ were harvested from 0.5-1.1 mm anterior to the bregma.

### Immunohistochemistry

Brain sections were washed 4×10 min in TBS at room temperature, followed by a 1 h blocking step in TBS^++^ (TBS with 0.3% horse serum (Millipore) and 0.3% Triton-X100 (Sigma)) at room temperature. The tissue was transferred to 0.5ml Safe Lock Reaction-Tubes containing 200 μl TBS^++^ including primary antibodies. Samples were incubated for 24-48 h at 4°C. Tissue samples were washed 4×10 min in TBS at room temperature, followed by a 30 min blocking step in TBS^++^ at room temperature. Brain sections were transferred to 0.5 ml Safe Lock Reaction-Tubes containing 200 μl TBS^++^ including secondary antibodies. Samples were incubated in the dark for 2 h at room temperature. Subsequently, brain slices were washed 4x 10 min in TBS at room temperature and were mounted on glass slides with Fluoromount G (eBioscience). The following antibodies were used: mouse anti-Sox2 (Abcam; 1:100), guinea pig anti-DCX (Merck; 1:400), rabbit anti-S100b (Abcam; 1:100), goat anti-mCherry (SICGEN; 1:1000) and chicken anti-GFAP (GeneTex; 1:500). Nuclei were counterstained with Hoechst 33342 (Biotrend; 1:3000).

### Microscopy and Cell Quantification

All images were acquired with a Leica TCS SP5 AOBS confocal microscope equipped with a UV diode 405 nm laser, an argon multiline (458-514 nm) laser, a helium-neon 561 nm laser, and a helium-neon 633 nm laser. Images were acquired as multichannel confocal stacks (z-plane distance 3 μm) in 8-bit format by using a 20x or 40x oil immersion objective at a resolution of 1024×1024 and 200Hz. For quantification of the v-SVZ and total brain sections, tile scans of the whole ventricle or the whole coronal brain section were acquired with a total z-stack size of 25μm. To quantify the OB, tiles cans of the whole OB covering the tissue thickness were acquired. For stained cells from *in vitro* culture, 4-9 fields of view were imaged. For representative images (2048×2048 resolution, 100Hz), the maximum intensity of a variable number of z-planes was stacked to generate the final z-projections. Representative images were cropped, transformed to RGB color format, and assembled into figures with Inkscape (inkscape.org). For cell quantification, ImageJ (NIH) was used including the plug-in cell counter to navigate through the z-stacks. To quantify cells in the OB, the volume of the OB was calculated by multiplying the entire area of every OB section (including the glomerular layer; GLL) with the entire z-stack size. Then we converted μm^3^ to mm^3^. Finally, cell counts were given as cells/mm^3^ OB. To elucidate the labeling efficiency of the different AAV variants in the total v-SVZ (medial, dorsal, and lateral wall of the lateral ventricle), the cells were counted on 25 μm thick coronal sections and are given as cells per 25 μm section. Mainly NSCs located in the lateral wall of the ventricle generate OB neurons during homeostasis. Since a particular area of the lateral v-SVZ serves cells to a particular volume of the OB, cell numbers were counted for the mathematical modeling of the lateral v-SVZ only. The length of the lateral ventricular wall was measured in a coronal section and multiplied with the z-stack size (25 μm), to estimate the area of the lateral v-SVZ. Afterwards, cells in the lateral v-SVZ were counted and normalized to the lateral v-SVZ area. Data are given as cells per mm^2^.

### NGS-screening of Barcoded AAV Capsid Variants - Computational Analysis

NGS-samples were sequenced and demultiplexed by the DKFZ Genomics and Proteomics Core Facility using bcl2fastq 2.19.0.316. This resulted in two (paired-end) FASTQ-files per sample. Each FASTQ consists of reads resulting from the targeted barcode amplification and up to 50% PhiX DNA that was spiked in to increase library complexity.

Each AAV variant is associated with a unique 15-mer barcode sequence. To quantify the most successful AAV, we simply counted how often each barcode occurred in each FASTQ file, bearing in mind the following pitfalls:

1. barcode-sequences might occur outside of the amplicon by chance, e.g. in the PhiX genome
2. barcodes might have sequencing errors
3. barcodes occur on the forward and reverse strand

To circumvent issues 1 and 2, we opted for a strategy where we only count barcodes matching the expected amplicon structure. This was achieved with the following regex (regular expression; defines a text search pattern): (?< = [NGCAT]{33}TGCTC)[NGCAT]{15}(?=CAGGG[NGCAT]{45}). Variable 15-mers [NGCAT]{15} are only counted if they are flanked by the expected regions TGCTC and CAGGG. Furthermore, we enforce a minimum of 33 upstream nucleotides and 15 downstream nucleotides, in addition to the flanking regions, to only count 15-mers at the expected position. 15-mers matching this regex were extracted and counted with the standard GNU command-line tools grep, sort and uniq. 15-mers sequenced from the reverse strand were counted with an equivalent reverse complement regex and added to the forward counts.

### Assigning Barcodes to AAV Capsids

Raw 15-mer counts were further processed in R. Most observed 15-mers matched a known barcode exactly (library #1: 74%, library #3: 87%), which allowed us to assign them to a unique AAV variant. The remaining 15-mer counts were added to the counts of the closest known barcode, allowing for a maximum of two mismatches.

### Normalization

Each sequenced sample corresponds to one tube with up to 500 FACS-sorted cells. To downweigh samples with lower cell numbers, barcode counts were scaled by the respective number of FACS events (usually 500, Supplementary Table 2). Barcode counts of the same cell type and biological replicate (termed “sets”) were then summed. The AAV libraries used for transduction contain slightly unequal proportions of AAV variants, which means that some AAV variants may have an advantage due to increased starting concentration. To remedy this problem, barcode counts were further scaled by their abundance in the transduction library (as determined by ^50^), so that barcode counts corresponding to more frequent AAV capsids were decreased and vice versa.

To account for sequencing depth of the individual samples, normalized barcode counts were divided by the total number of valid barcodes in that sample, yielding normalized barcode proportions. A potential source of bias is that amplicons with different barcodes may have different RT-PCR efficiencies. A previous study ^49^ on ten barcoded AAV variants found no such bias, but nonetheless we evaluated one possible source of bias, barcode GC-content, in our own data. We found no significant association between barcode GC-content and mean barcode proportion across all samples in either library (Fig S2 l-m).

### Identification of Candidate AAVs With High Transduction Efficiency

To identify the most promising AAV variants, AAVs were ranked by the mean normalized barcode proportion within and across cell types (Figure 1d-j). AAV1_P5 and AAV9_A2 performed consistently well across replicates of both experiments and were selected for further validation.

### Mathematical Modeling

A detailed description on how the mathematical modeling was developed is given in the Supplementary Material section.

### Statistics

Statistical analyses were performed with R version 4.0.2 using one-way ANOVA followed by Tukey’s Honest Significant Difference (HSD) post-hoc test unless otherwise noted. Tukey’s HSD p-values were corrected for multiple testing with the Benjamini-Hochberg procedure. The homogeneity of variance assumption of ANOVA was assessed with Levene’s test and the normality assumption was assessed with the Shapiro-Wilk normality test. The respective p-values are indicated in the figure legends. Figures were plotted with the R package ggplot2 and SigmaPlot 12.5.

## Data and Code Availability

All sequencing data is available at the NCBI Gene Expression Omnibus (GEO) under the accession GSE145172.

All scripts used in the analysis are available at https://github.com/LKremer/AAV-screening.

## Acknowledgements

We thank Monika Langlotz and the ZMBH FACS Core Facility; the DKFZ High Throughput Sequencing Unit; the DKFZ Microscopy Core Facility; Ellen Wiedtke and the members of the Dirk Grimm laboratory for technical assistance; Stefanie Limpert for technical assistance and the members of the Martin-Villalba laboratory for critically reading the manuscript.

## Author Contributions

SD was involved in project and experimental design, performed experiments including *in vitro* and *in vivo* screens, *ex vivo* NSC transplantation, *in vitro* and *in vivo* validations. LPMK was responsible for the bioinformatics analysis of all *in vitro* and *in vivo* screens and sequencing experiments. SK and SC conducted the single cell RNA sequencing experiment. SC conducted the FACS quantification of cells transduced with lead-candidates. TS was responsible for the mathematical modeling of the *in vivo* data. JW provided the two AAV capsid libraries and contributed to experimental design. HA and AL helped in producing AAV vectors. AM-C contributed to the development of the mathematical model, interpretation of data, and revision of the manuscript. DG, SA, AM-C and AM-V supervised the project and wrote the manuscript. AM-V designed and coordinated the study. All authors have read and approved the final version of the manuscript.

