## Supplementary material for "*In vivo* high-throughput screening of novel adeno-associated viral capsids targeting adult neural stem cells in the subventricular zone": Supplemental information.pdf

### Mathematical Modeling of Labeling Dynamics with AAV1_P5 transduced v-SVZ cells

#### Contents

**1 Mathematical Model of Neurogenesis** 1

**2 Model of Labeling Dynamics** 2

**3 Fitting of Model to Microscopic Data** 5
3.1 Data . . . 5
3.2 Fitting . . . 6

**4 Fitting the model to AAV1P5-CRE single cell data** 10

**5 Fitting of Model to FACS Data** 12

**6 Conclusion** 21

### 1 Mathematical Model of Neurogenesis

We extend our previously established mathematical model from [1–3]. The model describes time evolution of active NSC, quiescent NSC and TAPs. The model considers the following processes:

- Quiescent stem cells are activated at the rate $r$. As demonstrated in [1] the activation rate depends on the age of the organism.

- Division of active stem cells occurs at the rate $p_{stem}$. Upon division a stem cell gives rise to two progeny.
- The probability that a progeny is a stem cell is $b$. It is referred to as self-renewal probability. With probability $(1-b)$ the progeny differentiates into a TAP.
- TAPs divide a finite number of times before they further differentiate.

The part of the model describing stem cell dynamics has been parameterized in [1]. The TAP dynamics have not been calibrated so far. We assume a TAP doubling time of 20.15 hours taken from [4]. Assuming that TAP differentiate after two divisions we obtain the best agreement of model simulations with data, see Figure 1. We then have the following model.

$$\begin{aligned}
\frac{d}{dt}qNSC =& -r(t)\cdot qNSC + 2\cdot b\cdot p_{stem} aNSC \\
\frac{d}{dt}aNSC =& r(t)\cdot qNSC - p_{stem}\cdot aNSC \\
\frac{d}{dt}TAP_0 =& -p_{TAP}\cdot TAP_0 + 2\cdot(1-b)\cdot p_{stem}\cdot aNSC \\
\frac{d}{dt}TAP_1 =& -p_{TAP}\cdot TAP_1 + 2\cdot p_{TAP}\cdot TAP_0 \\
r(t) =& r_{max}\exp(-\beta_r t)
\end{aligned} \tag{1}$$

As $qNSC(t)$ and $aNSC(t)$ we denote the amount of quiescent and active neural stem cells at time $t$. As $TAP_i(t)$, $i\in\{0,\ 1\}$ we denote the amount of TAPs that have performed $i$ divisions at time $t$. Namely, $aNSC$ give rise to $TAP_0$. If $TAP_0$ divide, the progeny belong to $TAP_1$, i.e., TAP that have performed one division. Progeny of $TAP_1$ are neuroblasts. For notational convenience we omit the argument $t$ and identify $qNSC(t)\equiv qNSC$, $aNSC(t)\equiv aNSC$, $TAP_i(t)\equiv TAP_i$. Proliferation rates of stem cells and TAP are denoted as $p_{stem}$ and $p_{TAP}$ respectively. By $b$ we denote the probability of stem cell self-renewal. It is the probability with which a progeny of a stem cell is again a stem cell [5–7]. We note that the initial condition for TAPs has practically no impact on the cell counts at ages larger than 1 month. All model parameters are summarized in Table 1.

#### 2 Model of Labeling Dynamics

We make the following assumptions

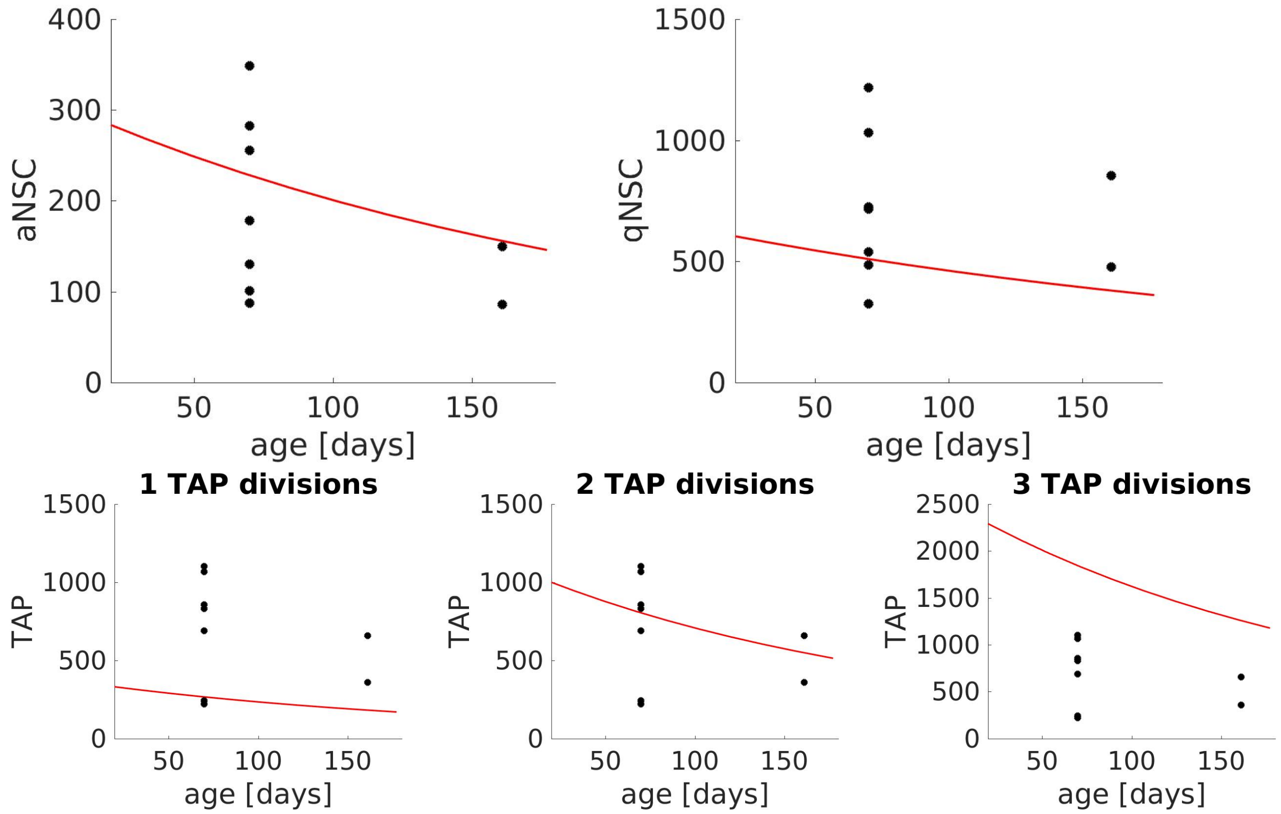

Figure 1: Simulation of NSC and TAP dynamics during aging. Upper: NSC dynamics. Lower: TAP dynamics assuming 1, 2 or 3 TAP divisions before further differentiation. We obtain the best agreement of model and data for 2 TAP divisions. Red curve: Model simulations, black dots: experimental data (FACS). The figures are obtained from simulation of model (1) with the parameters specified in Table 1.

| parameter | value |
|---|---|
| $r_{max}$ | $0.453\ d^{-1}$ |
| $\beta_r$ | $9.5 \cdot 10^{-4}\ d^{-1}$ |
| $b$ | 0.494 |
| $p_{stem}$ | $0.951\ d^{-1}$ |
| $p_{TAP}$ | $0.826\ d^{-1}$ |
| $qNSC(0)$ | 647 |
| aNSC(0) | 310 |

Table 1: Parameters of the neurogenesis model. Parameters are taken from [1].

- The transduced cell expresses a fluorescent label that is transmitted over the cell division. We assume that proliferation rates and self-renewal probability are not affected by the virus.
- The probability for a single TAP to be labeled is the same for $TAP_0$ and $TAP_1$.
- After the second division a TAP gives rise to two neuroblasts.
- Neuroblasts arrive after a delay $\theta$ in the olfactory bulb. During migration neuroblasts (NB) can divide and die. We model this by a sacling factor $\mu$. If proliferation outweights death $\mu > 1$, otherwise $\mu < 1$. The factor $\mu$ also takes into account the different volumes of SVZ and OB.
- We neglect the death of labeled OB cells during the duration of the experiment.

Dynamics of labeled cells is given by the system of equations (1) supplemented by the following equation for olfactory bulb neurons

$$\frac{d}{dt}OB = \mu \cdot 2 \cdot p_{TAP} \cdot TAP_1(t-\theta) \tag{2}$$

and the initial condition

$$\begin{aligned} qNSC(0) &= qNSC_0 \\ aNSC(0) &= aNSC_0 \\ TAP_i(0) &= TAP_{i,0},\ i \in \{0,\ 1\} \\ OB(0) &= 0. \end{aligned} \tag{3}$$

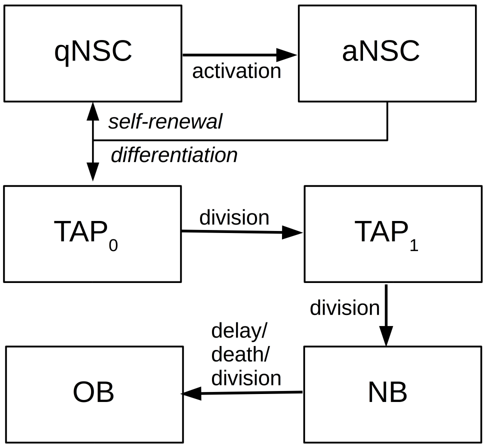

Figure 2: Model of serotype labeling. The scheme depicts the processes described by system (1)-(4). $qNSC$: labeled quiescent NSC, $aNSC$: labeled active NSC, $TAP_i$: labeled TAPs that have performed $i$ divisions, $NB$ : labeled neuroblasts, $OB$: labeled cells in the olfactory bulb.

The age of the mice at the beginning of the experiment is $\tau$. Since we define the time when the experiment starts as $t = 0$, the equation for $r$ is given by

$$r(t) = r_{max} \exp(-\beta_r(t + \tau)). \tag{4}$$

The model is visualized in Figure 2. In agreement with the quasi-steady state cell counts we set $TAP_{1,0} = 2 \cdot TAP_{0,0}$. For $t < 0$ all populations equal 0.

In the model we neglect the time between transduction and label expression. Shortly after the beginning of the experiment the number of experimentally detected labeled cells may differ from the labeled cell counts predicted by the model, since the cells transduced at day zero and counted as labelled cells in the model may not yet express sufficient label concentrations to be detected.

### 3 Fitting of Model to Microscopic Data

#### 3.1 Data

We consider densities of labeled NSC (given per $mm^2$ of SVZ) and labeled olfactory bulb neurons (given per $mm^3$ of olfactory bulb). The data shows high heterogeneity among individual mice, with more than an order of magnitude between individual measurements.

Instead of fitting the model to average cell counts we subdivide the data into two groups. The data and the subgroups are shown in Figure 3. We ask whether we can fit the data of both groups assuming that they differ only with respect to the number of initially labeled NSC and TAP.

Taking into account the heterogeneity among mice the data was assigned to two different groups as follows:

- The data at day 35 was subdivided into one group of mice showing high numbers of labeled cells (group 1) and one group of mice showing low numbers of labeled cells (group 2).
- The mice studied at day 7 show less heterogeneity, therefore they are not subdivided.
- Taking into account that stem cell numbers practically do not vary in a time interval of 28 days [1], we assign the data points acquired at day 7 to group 1. Assignment of this data to group 2 leads to a worse fit.

#### 3.2 Fitting

We use densities of labeled primitive cells and labeled of OB cells.

We assume:

- Counts of labeled primitive cells in the SVZ correspond to the sum of labeled aNSC and labeled qNSC.
- Labelled TAPs exist in the model but they do not contribute to the experimentally counted labeled primitive cells in the SVZ.

We assume that $r_{max}$, $\beta_r$, $b$, $p_{stem}$ and $p_{TAP}$ are not affected by the labeling and assume the values given in Table 1. These values are taken from [1]. We assume that $\mu$ and $\theta$ do not vary between group one and group two. The number of initially labeled cells, $qNSC_0$, $aNSC_0$, $\sum_{i=0}^{1} TAP_{i,0}$ may be different for both groups. We estimate the unknown parameters using fmincon from MATLAB (The MatWorks, Natic, USA). The cost functional passed to fmincon is a weighted least square functional with the inverse of the standard deviations as weights. We assume that the standard deviation of the labeled NSC count in

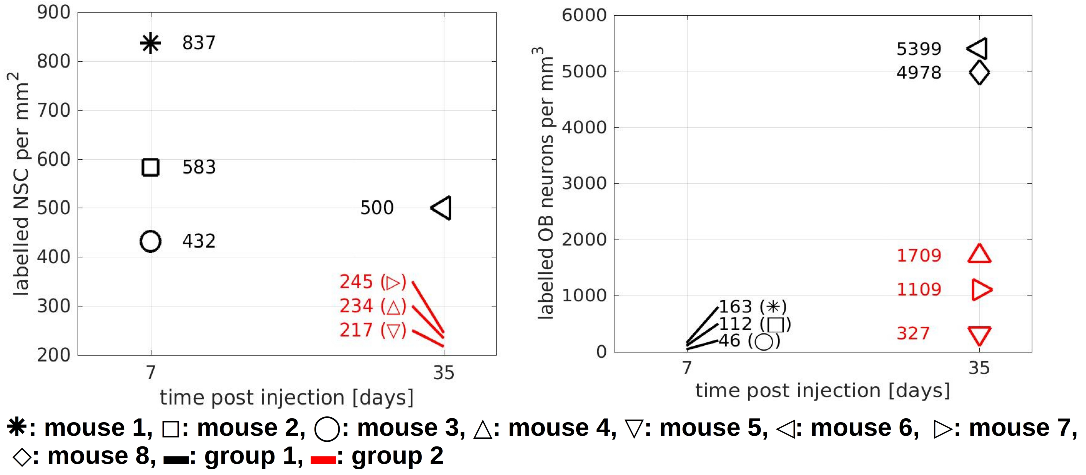

Figure 3: Time dynamics of labeled cells. Each mouse is identified by one symbol. Due to the heterogeneity among individual mice, each mouse was assigned to one of two groups. The color of the symbols indicates to which group the respective mouse belongs.

group 1 at 35 days is equal to the standard deviation of the labeled NSC count in group 2 at 35 days. We choose a multi-start approach with Latin hypercube sampling.

Our stem cell data is given per $mm^2$ of SVZ. Since there exist around 1000 NSC per $mm^2$ [8], we set an upper bound of 1000 for the initial number of labeled NSC.

The data from [1] imply that there exist approximately 5000 TAPs per 1000 NSC (standard deviation 1000) at the age where the experiments start. For the fitting we set an upper bound of 7000 TAPs per 1000 NSC.

Fitting different version of the model to the data implies that it has no impact on the estimated initial cell counts whether aNSC or qNSC are targeted by the virus (see next sections for details). Therefore, we assume in the following that active and quiescent NSC are labeled with the same probability.

The fitted parameters are provided in Table 2. The fit is depicted in Figure 4. Assuming that the labeling does not affect cell kinetics and that the observed heterogeneity comes from different numbers of initially labeled NSC and TAP, we obtain that in group 1 approx. 57%

| parameter | value |
|---|---|
| $aNSC_0 + qNSC_0$ | group 1: 569.2, group 2: 263.3 |
| $\sum_{i=0}^{1} TAP_{i,0}$ | group 1: 7000, group 2: 0.33 |
| $\theta$ | 6.9d |
| $\mu$ | 0.15m |

Table 2: Parameters obtained from the fit of the model to the experimental data. The model was fit to data from both groups simultaneously Only the number of initially labeled cells was allowed to be different for both groups. Other parameters are taken from [1].

of the NSC are labeled. According to the model in [1] at an age of 56 days approx. 69% of the NSC are quiescent and 31% are active. This corresponds to approximately 393 qNSC per $mm^2$ and 177 aNSC per $mm^2$ in group 1. In group 2 approximately 26% of NSC are labeled. This corresponds to approximately 179 qNSC per $mm^2$ and 81 aNSC per $mm^2$.

According to the parameterized model mice of age 56 days have 775 NSC. This implies that in group 1 442 cells were labelled initially and in group 2 202 cells were labelled initially.

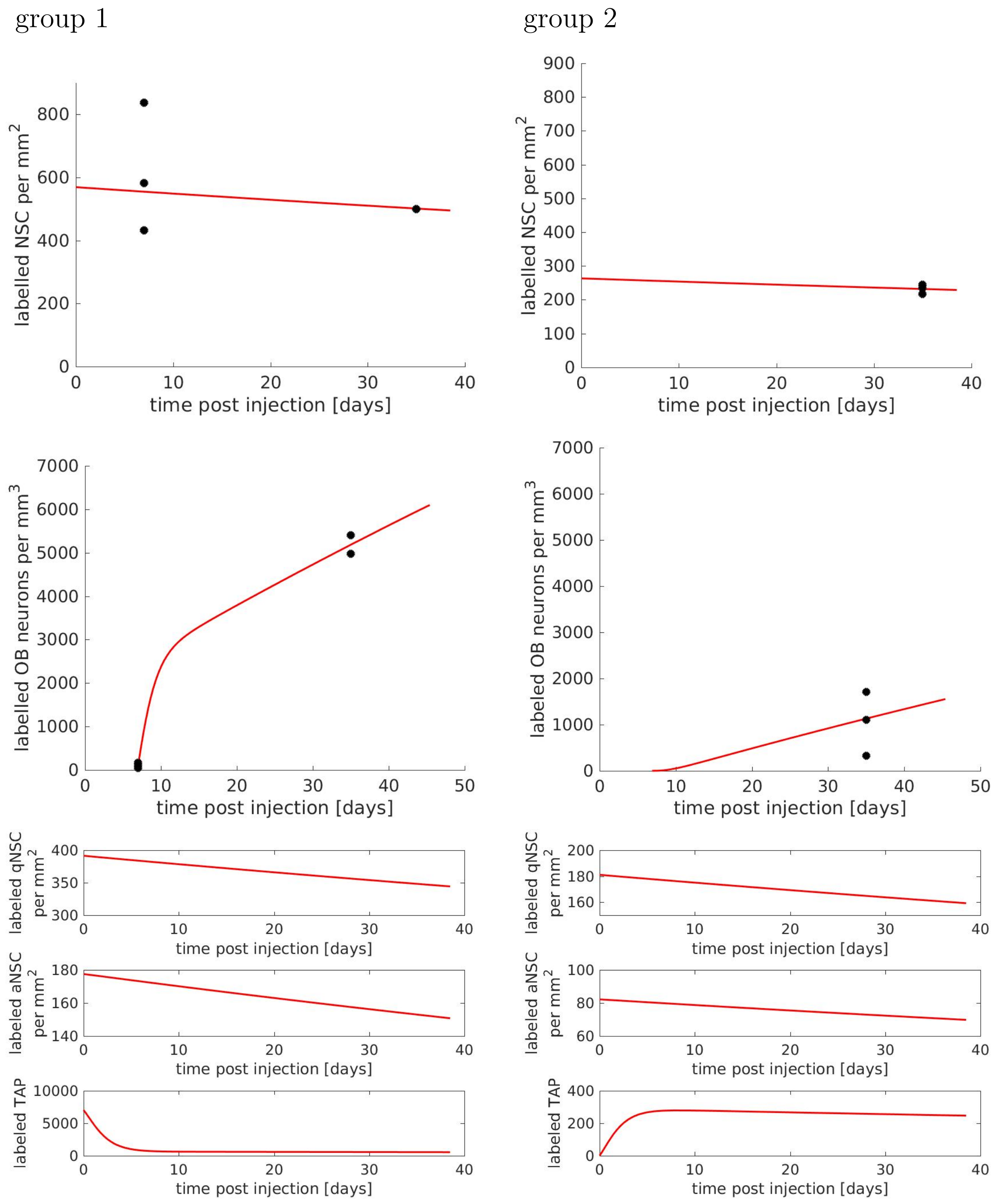

Figure 4: Comparison of model fit and data. The left column compares the fit to data from group 1, the right column to data from group 2. The model was fit to both groups simultaneously. Only the number of initially labeled NSC and TAP differs between the groups.

### 4 Fitting the model to AAV1P5-CRE single cell data

We validate the number of initially labelled NSC predicted by the model using single cell sequencing data.

The age of injection was $\tau = 91$ days. Cells were analyzed 37 days after injection. Due to the low numbers of sampled qNSC and the potentially low number of transcripts in quiescent cells, we we fit the model to TAP and aNSC data only.

To fit the model, we convert the cell fractions to absolute cell numbers. This is done by multiplying the fractions with the total cell counts for aNSC, qNSC and TAP obtained from the parameterized model for mice of age 91 days. In the model qNSC correspond to quiescent and resting cells, aNSC to actively cycling NSC. Approx. 50% of the aNSC counted in experiments do not cycle actively. For the modelling we consider these cells as qNSC.

As unknown parameters we consider the number of initially labeled $aNSC$, $qNSC$, $TAP$. We estimate the unknown parameters using fmincon from MATLAB (The MatWorks, Natic, USA). The cost functional passed to fmincon is a weighted least square functional with the inverse of the standard deviations as weights. We choose a multi-start approach with Latin hypercube sampling.

We consider two extreme scenarios, in the first scenario only aNSC and TAP are initially labelled (Fig. 5 ), in the second scenario only qNSC and TAP are initially labelled (Fig. 6)

We observe that both scenarios fit the data equally well and lead to the same value of the Akaike information criterion ($AIC_c$). The biological reason for this observation is that labelled quiescent cells become activated over time and labelled active cells can return to quiescence. Therefore, over time the labelled cells are distributed over both compartments, independently of their state at the time of labeling.

The number of initially labelled NSC of approximately 620 cells is similar to the labelled stem cell counts inferred from model-fitting to microscopic data (440 cells for the highly labelled group).

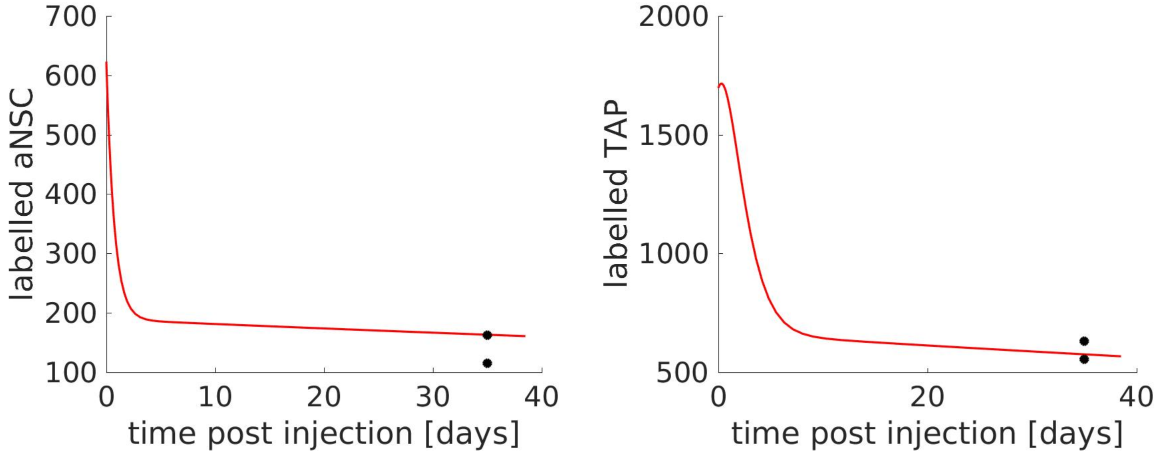

Figure 5: AAV1P5-CRE single cell (only aNSC targeted). Estimated parameters: $qNSC(0) = 0$, $aNSC(0) = 623$, $TAP(0) = 1696$, $AIC_c = 15.1$

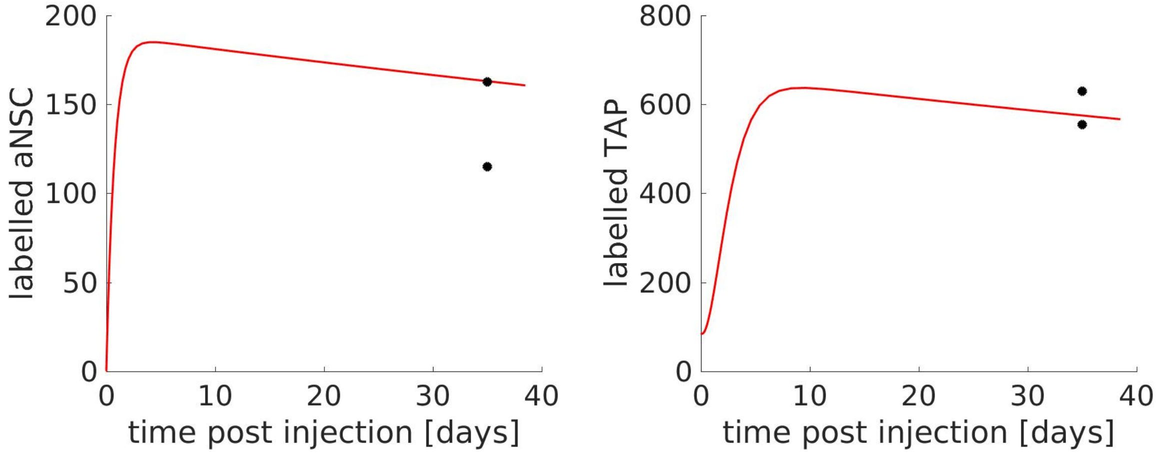

Figure 6: AAV1P5-CRE single cell (only qNSC targeted). Estimated parameters: $qNSC(0) = 617$, $aNSC(0) = 0$, $TAP(0) = 85$, $AIC_c = 15.1$

### 5 Fitting of Model to FACS Data

The microscopic data cannot distinguish between quiescent and active NSC. For this reason we consider FACS data. We use the FACS data to investigate labelled cell kinetics on short time scales. The serotype is injected in mice of age $\tau = 63$ days and $\tau = 158$ days respectively. We denote the time of serotype injection as $t = 0$.

We model the dynamics of labelled calls. Labelled aNSC are identified by their Glast+ / Prom+ / EGFR+ / YFP+ phenotype, labelled qNSC by their Glast+ / Prom+ / EGFR- / YFP+ phenotype. In the model $aNSC$ correspond to actively cycling cells and qNSC to quiescent and resting cells. Since 50% of the Glast+ / Prom+ / EGFR+ / YFP+ cell fraction are Ki67+, we fit the aNSC of the model to 50% of the experimentally obtained Glast+ / Prom+ / EGFR+ / YFP+ cell counts. The qNSC of the model are fitted to the experimentally counted Glast+ / Prom+ / EGFR- / YFP+ cell count plus the remaining 50% of the experimentally counted Glast+ / Prom+ / EGFR+ / YFP+ cells, which are Ki67-.

As unknown parameters we consider the number of initially labeled $aNSC$, $qNSC$, $TAP$, the delay $\theta$ and the amplification factor $\mu$. We estimate the unknown parameters using fmincon from MATLAB (The MatWorks, Natic, USA). The cost functional passed to fmincon is a weighted least square functional with the inverse of the standard deviations as weights. We choose a multi-start approach with Latin hypercube sampling.

#### 5.1 AAV1P5-YFP

The injection took place at the age $\tau = 63$ days. The cells were counted 6 days after injection. We test two extreme scenarios, in the first scenario only aNSC are initially labelled (Fig. 7), in the second scenario only qNSC are initially labelled (Fig. 8)

We observe that both scenarios fit the data equally well and lead to the same value of the Akaike information criterion ($AIC_c$). The biological reason for this observation is that labelled quiecent cells become activated over time and labelled active cells can return to quiescence. Therefore, over time the labelled cells are distributed over both compartments, independently of their state at the time of labeling. The model indicates that this redistribution takes place on a short time scale such that already 6 days after injection it cannot be distinguished if active or quiescent NSC were initialy targeted by the label.

This implies that the model does not allow to infer at which state cells have been labeled. However, the model implies that, independently of whether qNSC or aNSC are targeted by the virus, already 6 days after injection labelled cells exist in both NSC states.

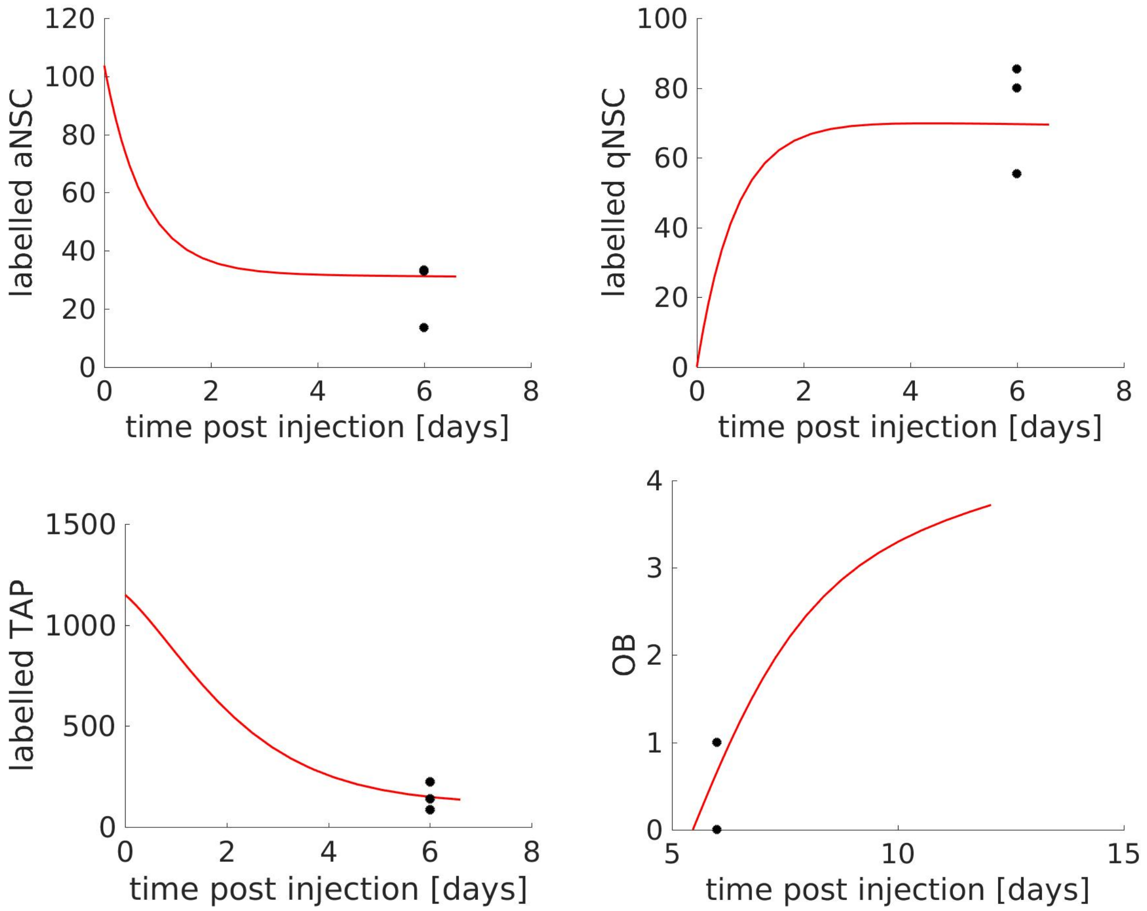

Figure 7: AAV1P5-YFP (only aNSC targeted). Estimated parameters: $aNSC(0) = 103$, $qNSC(0) = 0$, $TAP(0) = 1150$, $\theta = 5.5d$, $\mu = 0.001$, $AIC_c = 9.8$

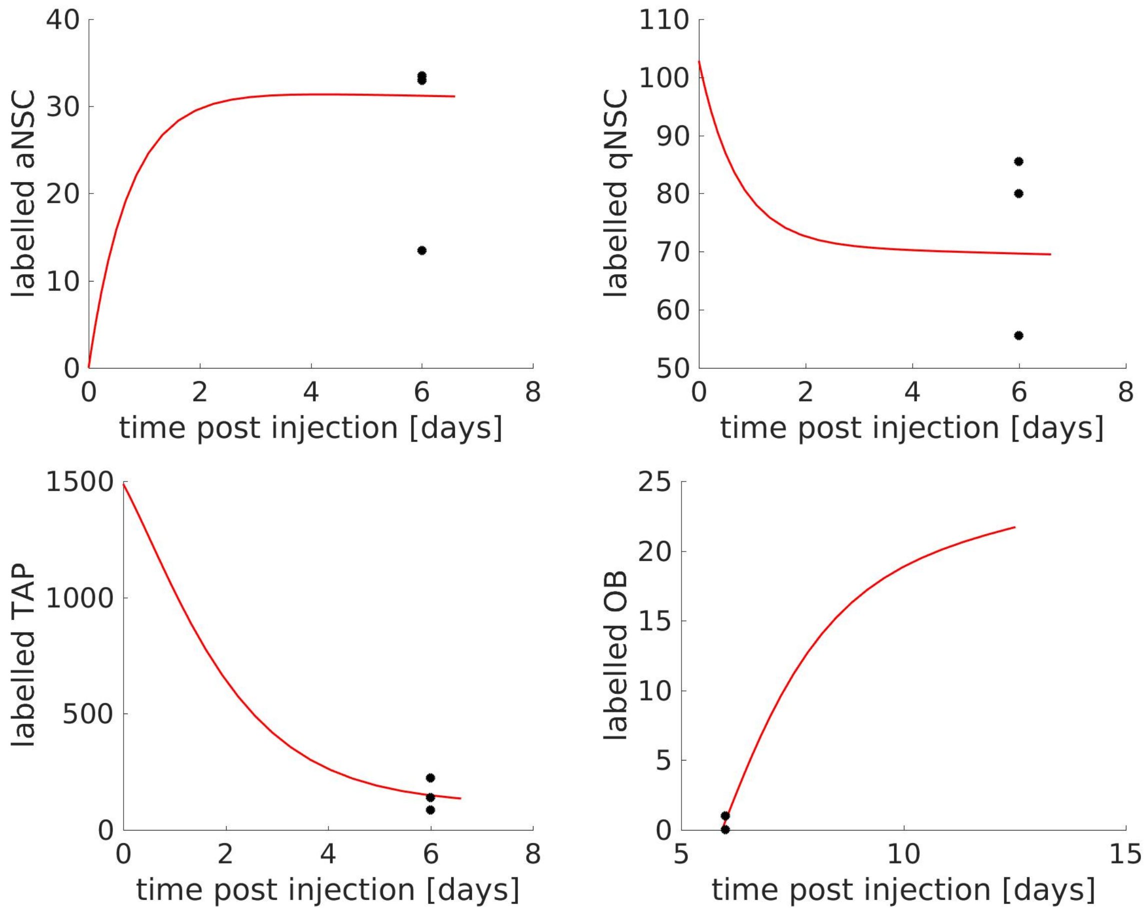

Figure 8: AAV1P5-YFP (only qNSC targeted). Estimated parameters: $qNSC(0) = 103$, $aNSC(0) = 0$, $TAP(0) = 1488$, $\theta = 5.9d$, $\mu = 0.005$, $AIC_c = 9.8$

##### 5.2 AAV9A2-YFP

The injection took place at the age $\tau = 63$ days. The cells were counted 6 days after injection. As before, we test two extreme scenarios, in the first scenario only aNSC and TAP are initially labelled (Fig. 9), in the second scenario only qNSC and TAP are initially labelled (Fig. 10)

As for the dataset considered before, it makes no difference for the labelled cell counts at day 6 whether we initially label only active or only quiescent cells, or an arbitarty mixture of them.

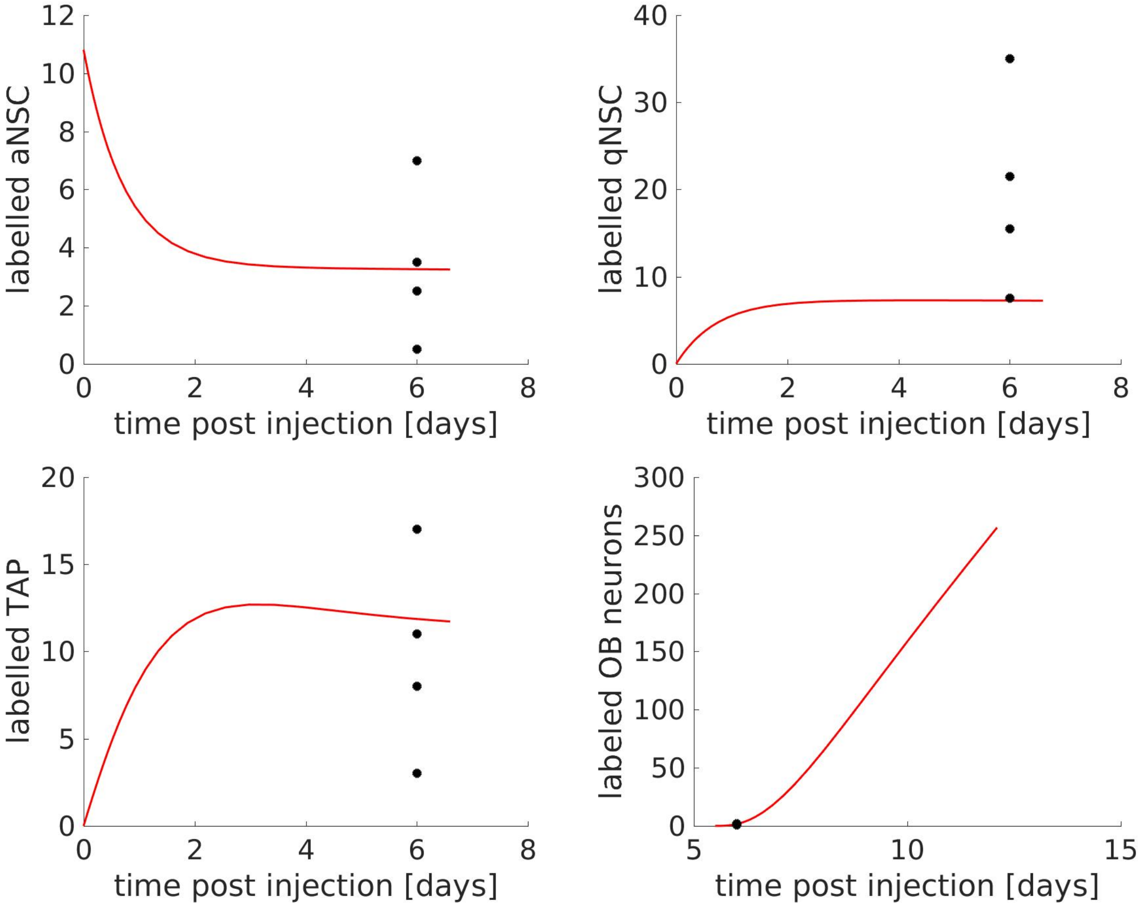

Figure 9: AAV9A2-YFP (only aNSC targeted). Estimated parameters:: $aNSC(0) = 11$, $qNSC(0) = 0$, $TAP(0) = 0$, $\theta = 5.5d$, $\mu = 3.5$, $AIC_c = 12.8$

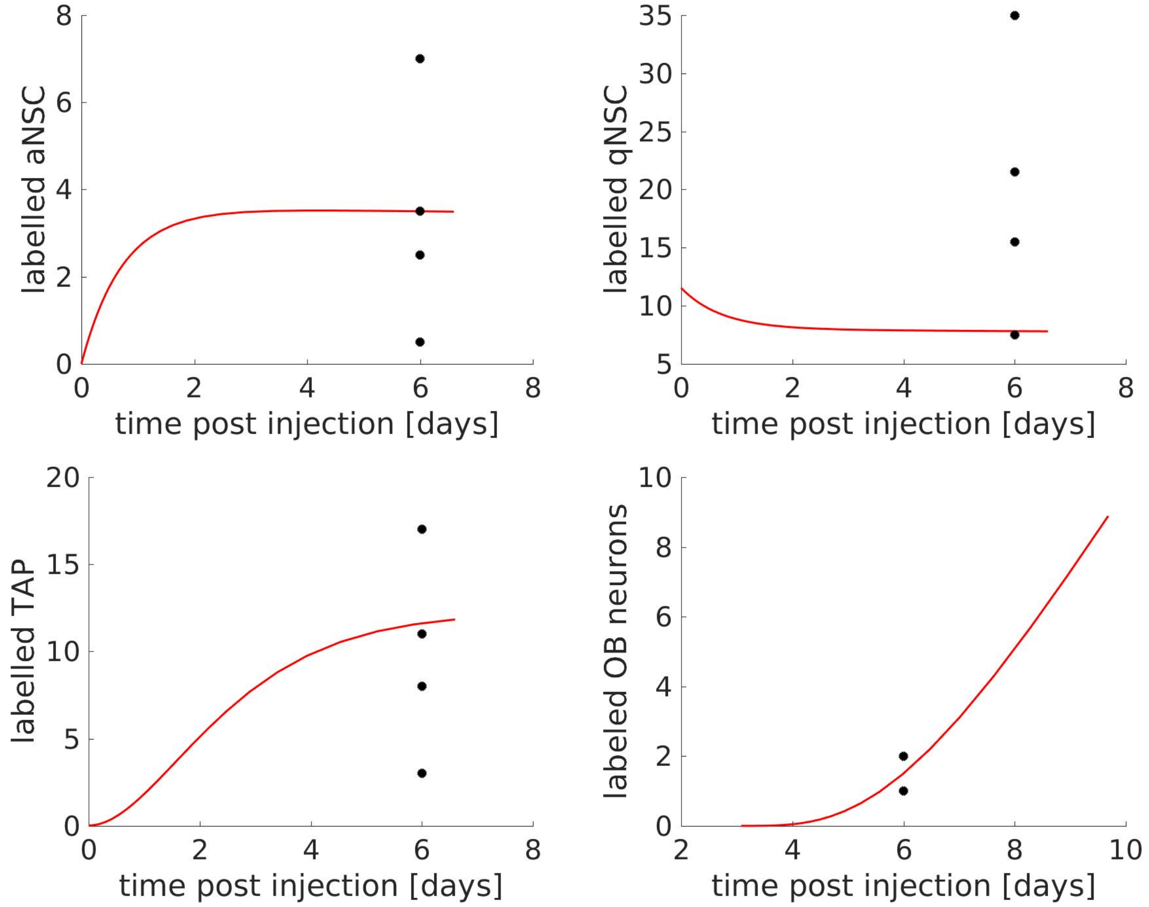

Figure 10: AAV9A2-YFP (only qNSC targeted). Estimated parameters: $qNSC(0) = 11$, $aNSC(0) = 0$, $TAP(0) = 0$, $\theta = 3.1d$, $\mu = 0.18$, $AIC_c = 12.4$

#### 5.3 AAV1P5-CRE

The injection took place at the age $\tau = 158$ days. The cells were counted 8 days after injection. We again test two extreme scenarios, in the first scenario only aNSC and TAP are initially labelled (Fig. 11), in the second scenario only qNSC and TAP are initially labelled (Fig. 12)

As for the dataset considered before, it makes no difference whether we label only active or only quiescent cells, or an arbitarty mixture of them.

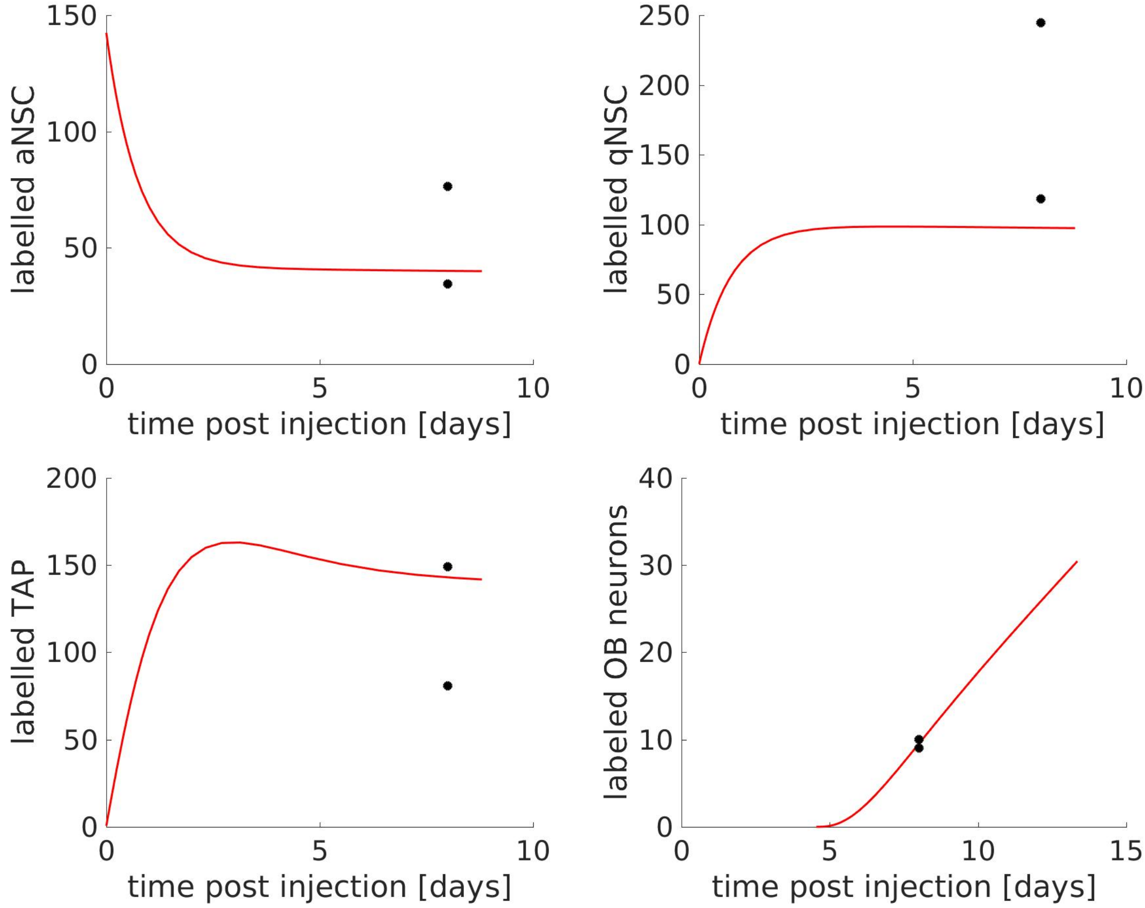

Figure 11: AAV1P5-CRE (only aNSC targeted). Estimated parameters: $qNSC(0) = 0$, $aNSC(0) = 142$, $TAP(0) = 1$, $\theta = 4.54d$, $\mu = 0.024$, $AIC_c = 20.3$

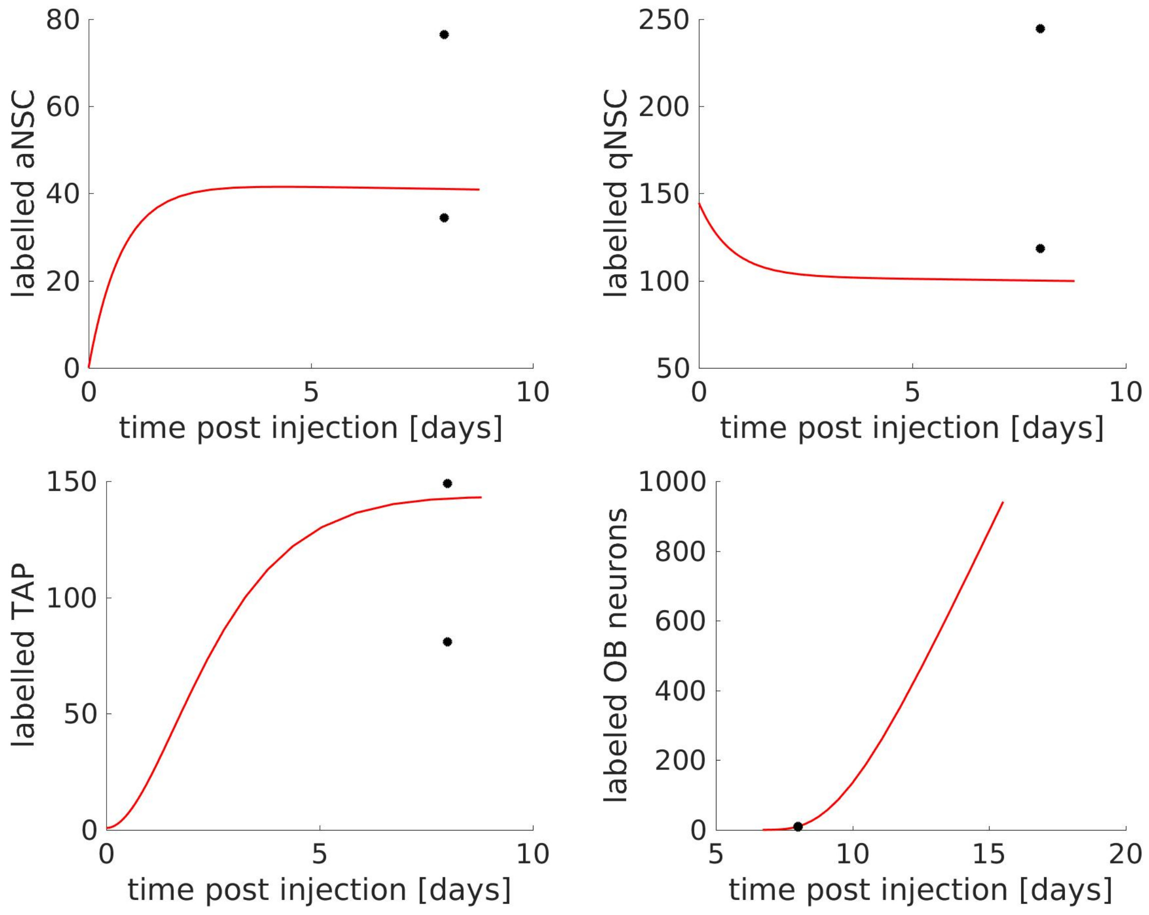

Figure 12: AAV1P5-CRE (only qNSC targeted). Estimated parameters: $qNSC(0) = 145$, $aNSC(0) = 0$, $TAP(0) = 1$, $\theta = 6.7d$, $\mu = 1.03$, $AIC_c = 20.0$

#### 5.4 Joined fit to AAV1P5-YFP, AAV1P5-CRE and AAV9A2-YFP

Assuming that $\theta$ and $\mu$ do not change on the considered age scale, we simultaneously fit the model to the AAV1P5-YFP, AAV9A2-YFP and AAV1P5-CRE datasets. For each dataset $\theta$ and $\mu$ are considered to be identical. The number of initially labelled cells can be different for each dataset.

The results are displayed in Fig. 13 and 14.

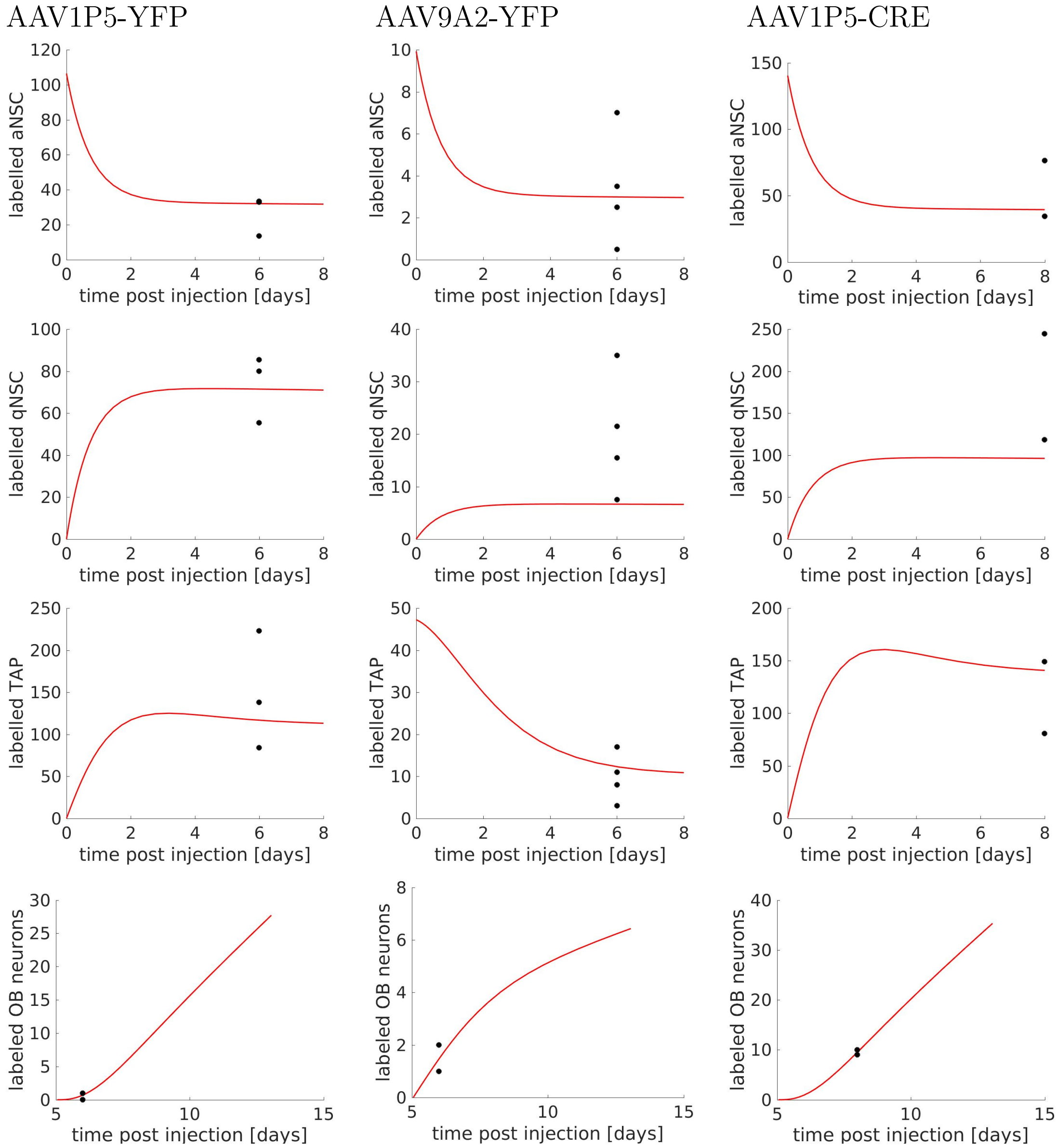

Figure 13: AAV1P5-YFP, AAV9A2-YFP and AAV1P5-CRE (only aNSC targeted). First column (AAV1P5-YFP) estimated parameters: $aNSC(0) = 106$, $qNSC(0) = 0$, $TAP(0) = 0$. Second column (AAV2P9-YFP) estimated parameters: $aNSC(0) = 10$, $qNSC(0) = 0$, $TAP(0) = 47$. Third column (AAV1P5-CRE) estimated parameters: $aNSC(0) = 140$, $qNSC(0) = 0$, $TAP(0) = 0$. For all datasets: $\mu = 0.03$, $\theta = 5.1d$, $AIC_c = 19.7$

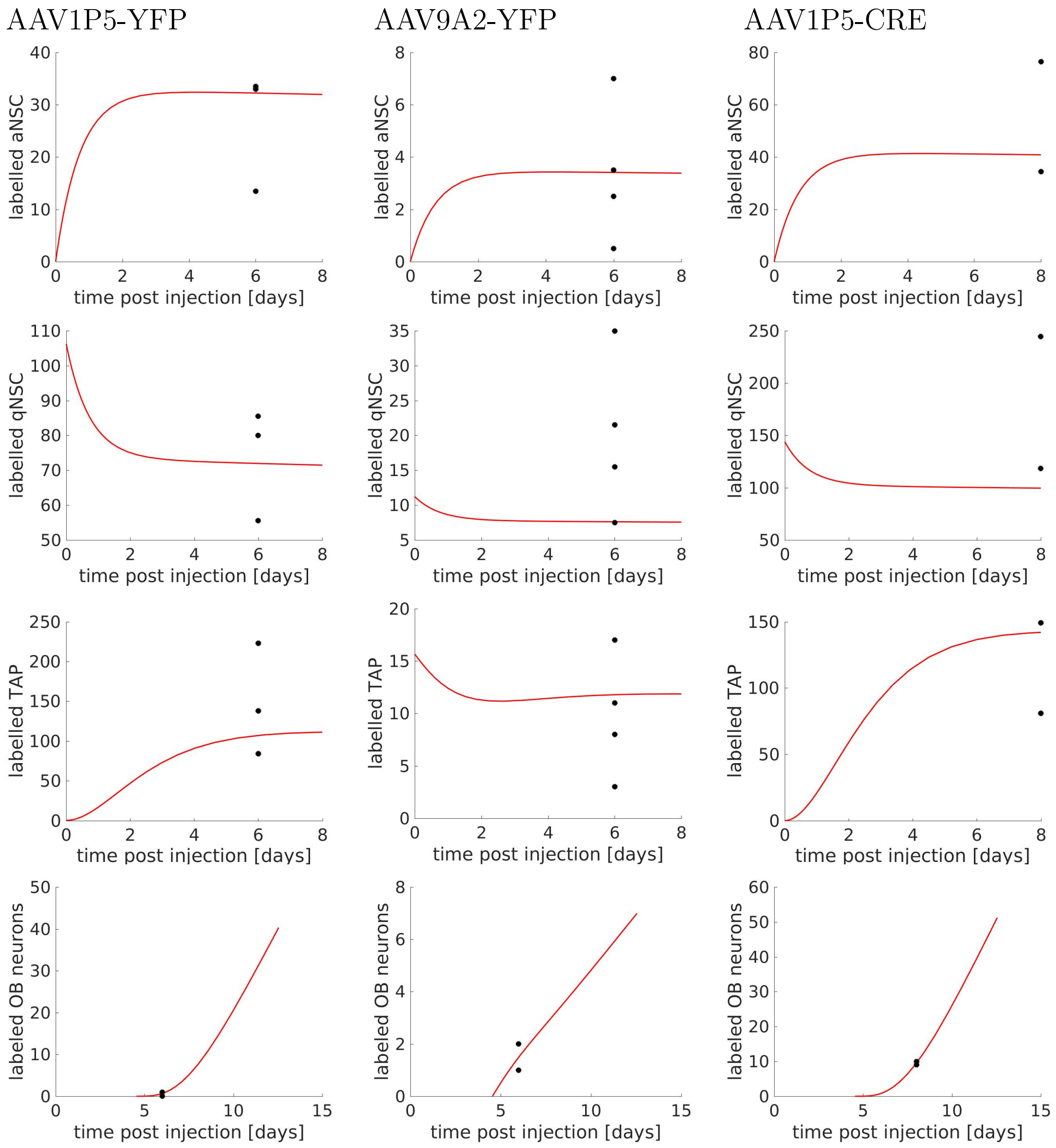

Figure 14: AAV1P5-YFP, AAV9A2-YFP and AAV1P5-CRE (only qNSC targeted). First column (AAV1P5-YFP) estimated parameters: $aNSC(0) = 0$, $qNSC(0) = 106$, $TAP(0) = 0$. Second column (AAV2A9-YFP) estimated parameters: $aNSC(0) = 0$, $qNSC(0) = 11$, $TAP(0) = 16$. Third column (AAV1P5-CRE) estimated parameters: $aNSC(0) = 0$, $qNSC(0) = 144$, $TAP(0) = 0$. For all datasets: $\mu = 0.066$, $\theta = 4.5d$, $AIC_c = 18.8$

### 6 Conclusion

In summary, the model implies that the measured labelled cell counts are in agreement with the hypothesis that cell properties do not change due to the adenovirus. The labeling efficiency inferred from the microscopic data with high amounts of labelled cells is similar to the labelling efficiency inferred from the single cell data. The ratio of labelled aNSC and labelled qNSC 6 days after virus injection is independent of the state of the NSC targeted by the virus.

### Supplementary Figures

#### Supplementary Figure S1

a

FACS-sorting strategy to isolate qNSCs; aNSCs; TAPs and neuroblasts

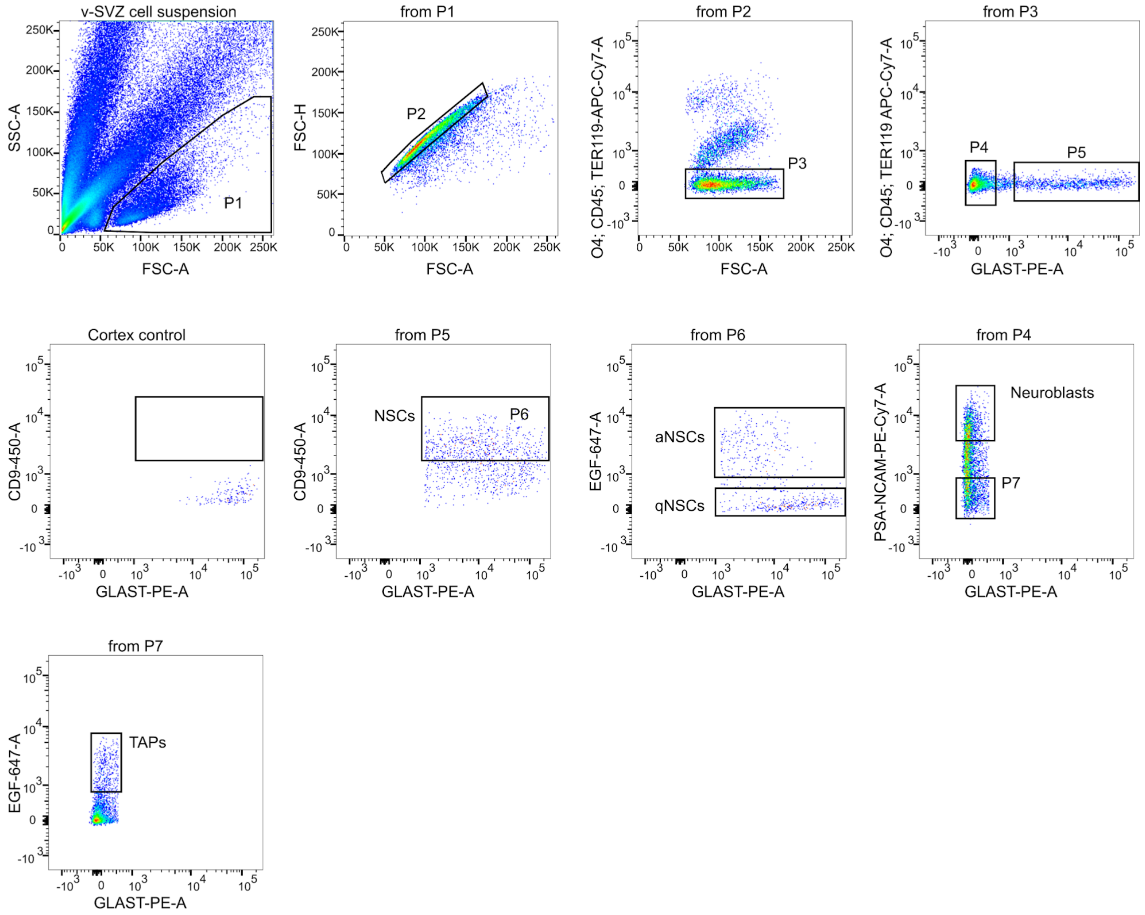

b

FACS-sorting strategy to isolate oligodendrocytes; astrocytes and ependymal cells

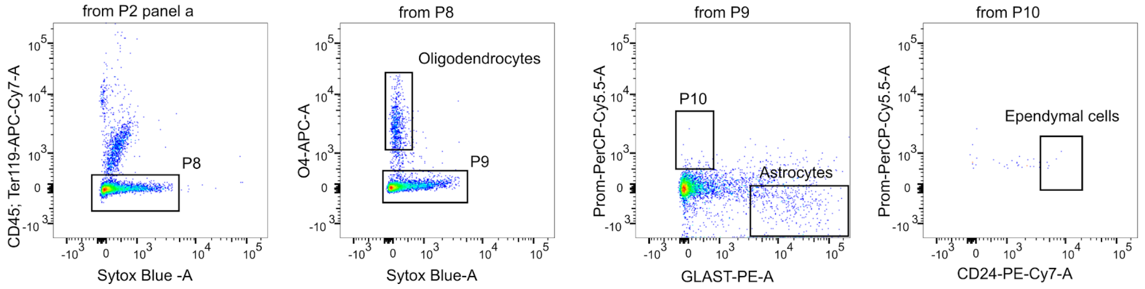

#### Figure S1, related to Figure 1:

**a** FACS sorting strategy to isolate total NSCs, qNSCs, aNSCs, TAPs and neuroblasts. **b** FACS sorting strategy to isolate oligodendrocytes, astrocytes and ependymal cells.

#### Supplementary Figure S2

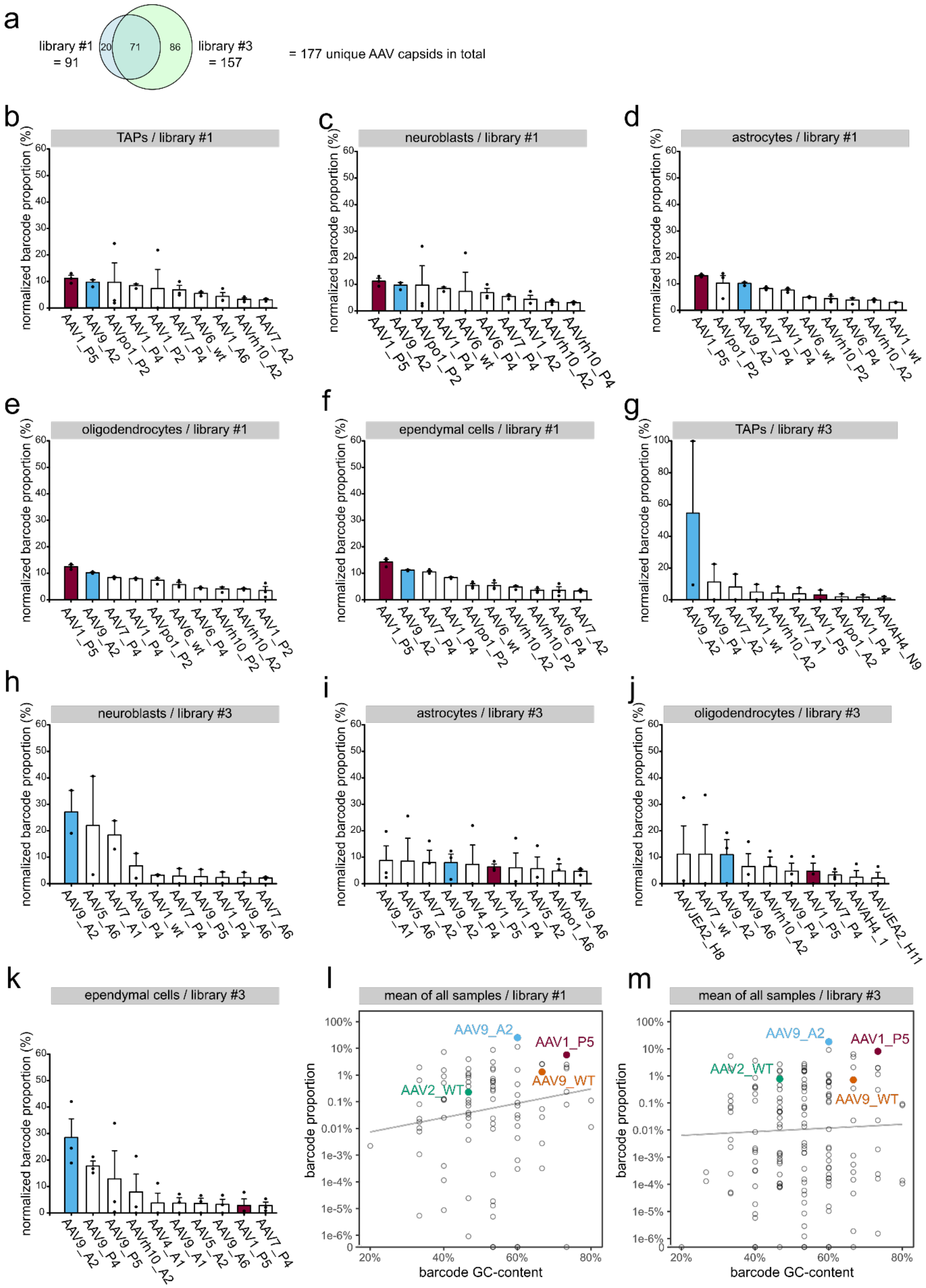

#### Figure S2, related to Figure 1:

**a** Number of barcoded AAV capsids in library #1 and library #3. 71 capsids are contained in both libraries. **b-f** Normalized barcode proportion over different FACS-sorted cell types seven days after library #1 transduction of **b** TAPs, **c** neuroblasts, **d** astrocytes, **e** oligodendrocytes and **f** ependymal cells. **g-k** Normalized barcode proportion over different FACS-sorted cell types seven days after library #3 transduction of **g** TAPs; n=2 sets, **h** neuroblasts; n=2 sets, **i** astrocytes, **j** oligodendrocytes and **k** ependymal cells. **l,m** Correlation of barcode GC-content and mean barcode proportion across all samples of library #1 (**l**; Spearman's $\rho=0.10$; $p=0.33$) and library #3 (**m**; Spearman's $\rho=0.07$; $p=0.36$). All mice were eight weeks old at the time of AAV injection, and all values are given as mean ± SEM; n=3 sets unless stated otherwise.

### Supplementary Figure S3

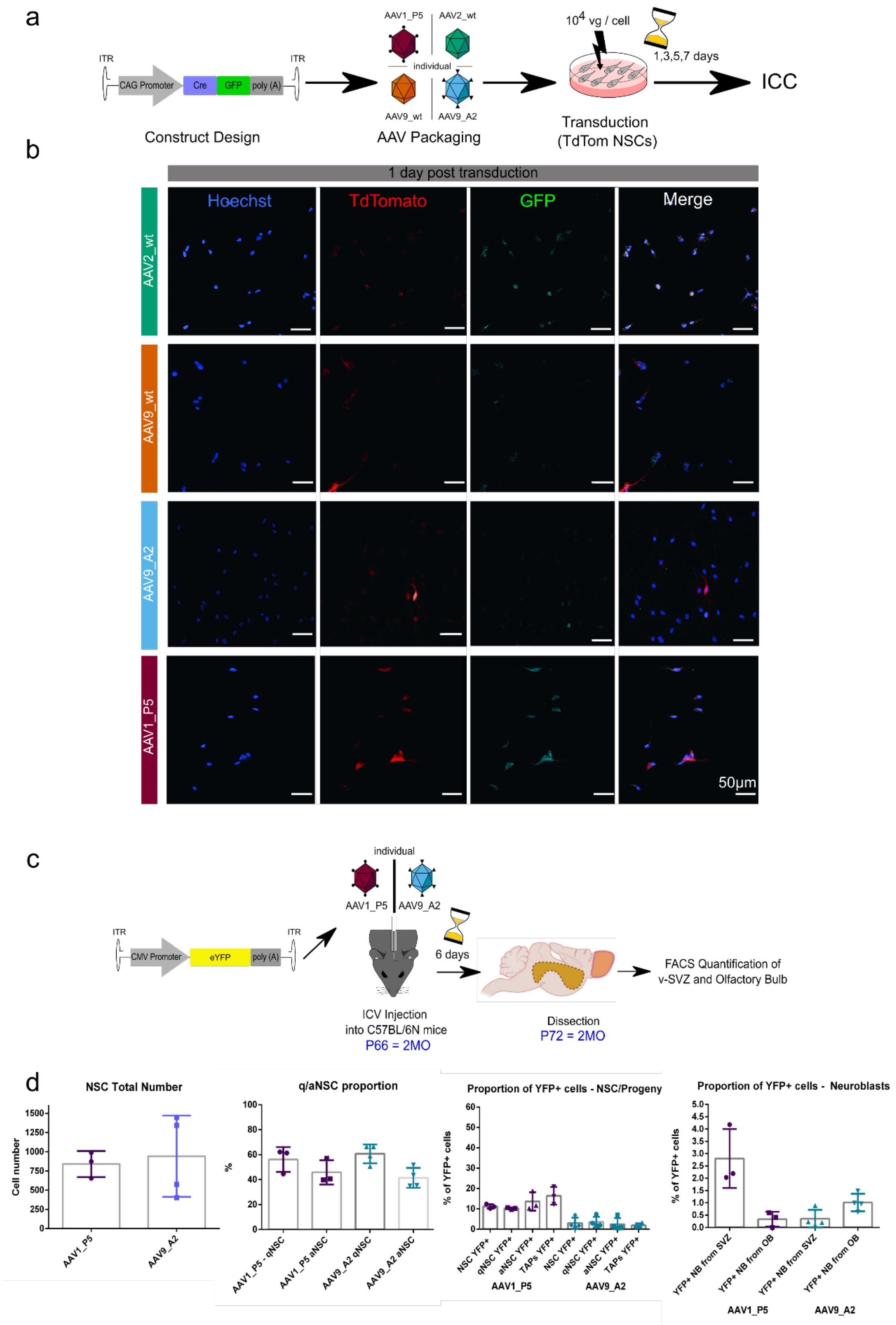

### Figure S3, related to Figure 2

**a** Schematic illustration of the experimental outline to *in vitro* validate different AAV capsids. **b** Representative images of NSCs *in vitro* transduced with different AAV capsids at day 1 after transduction; scale bar 50 µm. **c** Schematic illustration of the experimental outline to perform labeling efficiency analysis of the SVZ and olfactory bulb by FACS Quantification using either AAV1_P5_eYFP or AAV9_A2_eYFP ($10^{10}$vg/mouse). **d** Quantification of total NSC number in the v-SVZ; proportion of quiescent to active NSCs; labeling efficiency of NSC and TAPs in the v-SVZ; and labeling efficiency of neuroblasts in the v-SVZ and olfactory bulb. The overall NSC labeling efficiency was higher with AAV1_P5 (11.19% ± 0.63, n=3) compared to AAV9_A2 (2.96 ± 1.34, n=4) ($p<0.01$, two-sided Student's t-test). All values are given as mean ± SEM.

#### Supplementary Figure S4

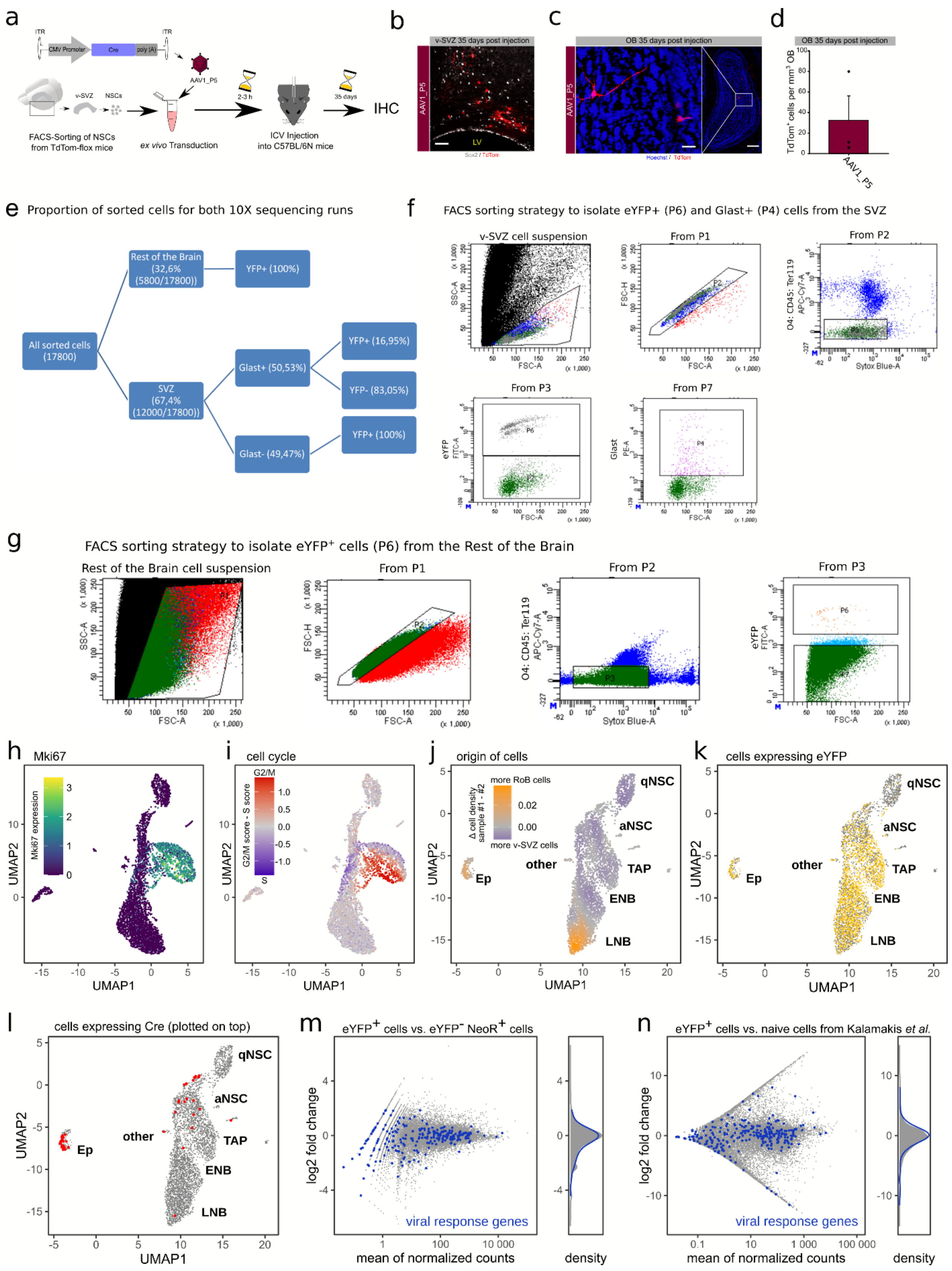

**Figure S4, related to Figure 3**

**a** Schematic illustration of the experimental outline to perform *ex vivo* manipulation and transplantation of NSCs. **b** IHC of the v-SVZ (scale bar 50 µm) and **c** Olfactory bulb (OB) neurons (scale bar 200 µm and 30 µm). **d** Quantification of tdTomato-positive cells in the OB; n=3. For this part, all mice were eight weeks old at the time of stereotactic injection of AAVs. **e** Average proportion of sorted cells from the v-SVZ and the rest of the brain (RoB, consisting of striatum, rostral migratory stream and OB) for both single cell RNA sequencing runs. The percentage of $eYFP^+$ and $GLAST^+$ cells from the sorted cells in every tissue is also shown. **f** FACS sorting strategy to isolate $eYFP^+$ and also $eYFP^-/GLAST^+$ cells from the v-SVZ. **g** FACS sorting strategy to isolate $eYFP^+$ cells from the RoB. **h-l** 2D representation of single cell transcriptomes before regressing out the effects of cell cycle heterogeneity (h,i) and after (j-l). **h** Cells expressing Mki67 (proliferation marker protein Ki-67, log-normalized UMI counts) form a distinct group. **i** Cell cycle phase scores highlight cells expressing canonical markers of S phase (blue) and G2/M phase (red). **j** Putative RoB cells are located at the end of the NSC lineage. Cell color indicates whether nearby cells mostly stem from sample #1 or sample #2 (see Methods for details). Sample #1 contains more cells from RoB, hence orange cells in the main lineage are mostly from RoB. **k,l** Cells with at least one eYFP (**k**) or Cre (**l**) transcript are highlighted in yellow or red. **m,n** MA plots of gene expression differences between $eYFP^+$ cells and $eYFP^-$ $NeoR^+$ cells (**m**) or $eYFP^+$ cells and untransduced cells from[2] (**n**). Right: $\log_2$ fold change distribution for all genes (gray) and viral response genes (blue). RoB, rest of the brain (entails the striatum, rostral migratory stream and olfactory bulb).

#### Supplementary Figure S5

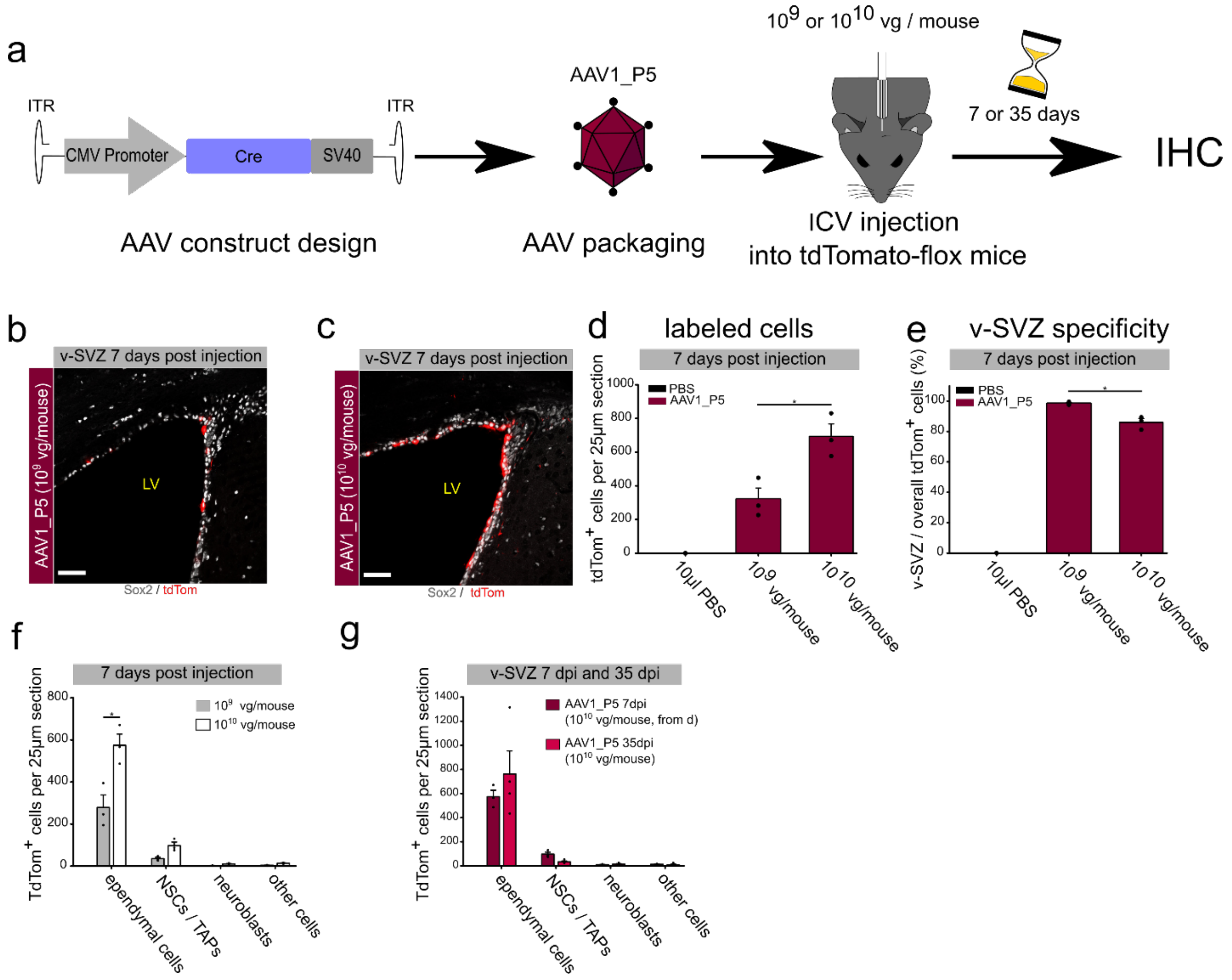

**Figure S5 related to Figure 4**

**a** Schematic illustration of the experimental outline to test v-SVZ labeling with different AAV concentrations. **b,c** IHC of the v-SVZ after injecting **b** $10^9$ or **c** $10^{10}$ vg per mouse (scale bar 50 µm). **d** Quantification of the total number of tdTomato-labeled cells within the v-SVZ injected with $10^9$ vg per mouse (319.89 ± 66.2) vs. $10^{10}$ vg per mouse (694 ± 73.92). **e** Quantification of tdTomato-labeled cells located in the v-SVZ among all tdTomato-positive cells in a 25µm thick coronal brain section ($10^9$ vg/mouse (98.4% ± 0.612) vs. $10^{10}$ vg/mouse (86.0 % ± 2.51)). **f** Quantification of ependymal cells ($10^9$ vg per mouse (278.67 ± 59.82) vs. $10^{10}$ vg per mouse (574.22 ± 53.74)), NSCs, neuroblasts and other cells in the v-SVZ. **g** Quantification of the total number of tdTomato-labeled cells within the v-SVZ of mice at 35dpi; n=4. All mice were eight weeks old at the time of AAV injection; n=3. All values are given as mean ± SEM; **$p \leq 0.01$ and ***$p \leq 0.001$ (Student's t-test). Cre, Cre recombinase; SV40, Simian-Virus 40 polyA signal; ICV, Intracerebroventricular; IHC, immunohistochemistry.

#### Supplementary Figure S6

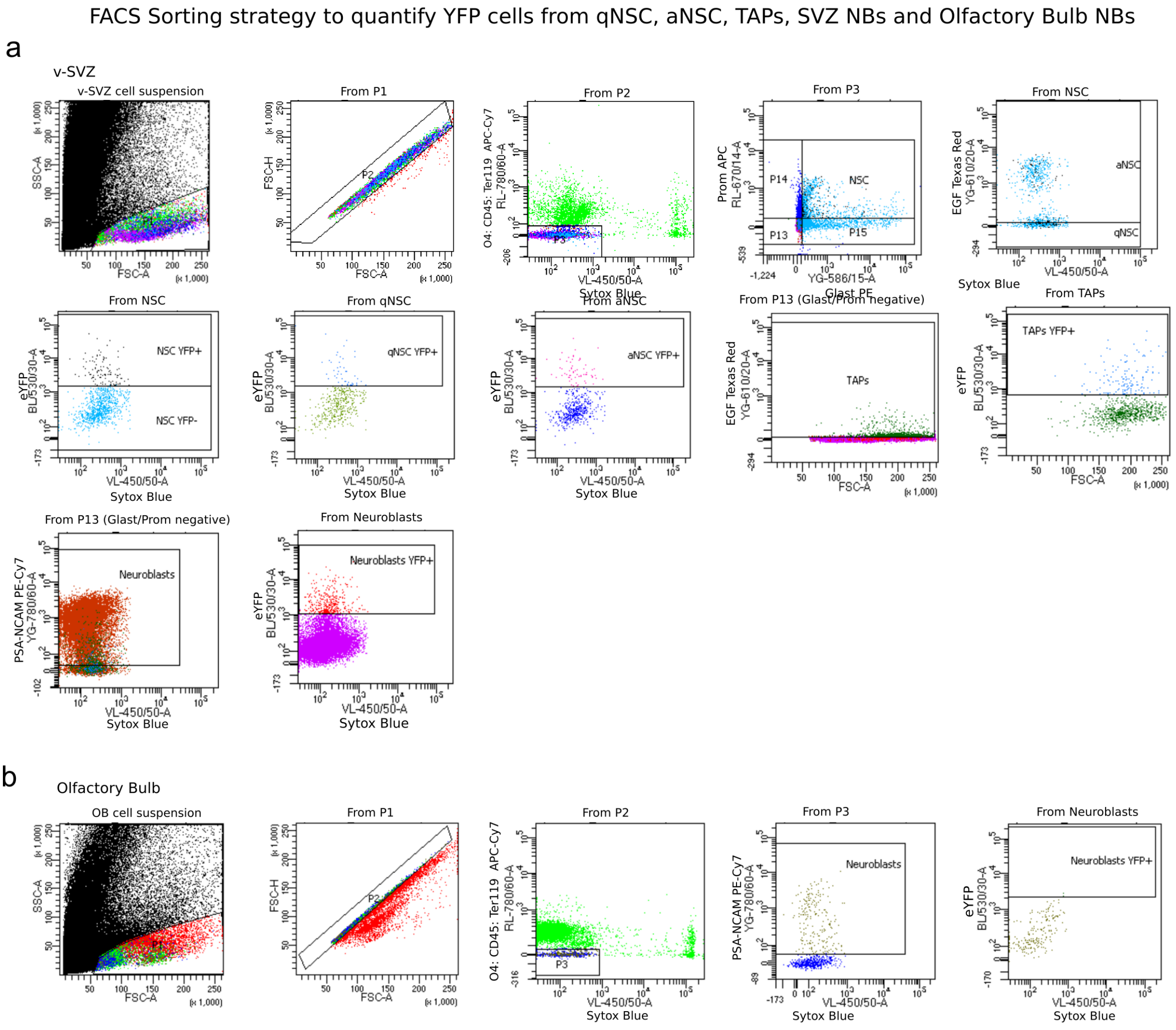

#### Figure S6 related to Figure 4

**a** FACS sorting strategy to quantify the percentage of YFP$^{+}$ cells from qNSCs, aNSCs, TAPs and NBs in the v-SVZ (related to Fig S3c-d and 4f-g). **b** FACS sorting strategy to quantify YFP$^{+}$ cells from NBs in the olfactory bulb.
